# Transcriptome sequencing and delimitation of cryptic *Oscarella* species (*O. carmela* and *O. pearsei* sp. nov) from California, USA

**DOI:** 10.1101/126326

**Authors:** Alexander Ereskovsky, Daniel J. Richter, Dennis V. Lavrov, Klaske J. Schippers, Scott A. Nichols

**Affiliations:** Institut Méditerranéen de Biodiversité et d’Ecologie Marine et Continentale (IMBE), CNRS, IRD, Aix Marseille Université, Avignon Université, Station Marine d’Endoume, 13007 Marseille, France; Department of Embryology, Faculty of Biology, Saint-Petersburg State University, 7/9 Universitetskaya emb., St. Petersburg 199034, Russia; Department of Molecular and Cell Biology, University of California, Berkeley, CA 94720-3200, USA; Sorbonne Universités, UPMC Université Paris 06, CNRS, UMR 7144, Adaptation et Diversité en Milieu Marin, Equipe EPEP, Station Biologique de Roscoff, 29680 Roscoff, France; Department of Ecology, Evolution, and Organismal Biology, Iowa State University, 2200 Osborn Drive, Ames, IA 50011, USA; Department of Biological Sciences, 2101 E. Wesley Ave., SGM 203, University of Denver, Denver, CO 80208, USA

## Abstract

The homoscleromorph sponge *Oscarella carmela*, first described from central California, USA is shown to represent two morphologically similar but phylogenetically distant species that are co-distributed. We here describe a new species as *Oscarella pearsei*, sp. nov. and redescribe *Oscarella carmela*; the original description was based upon material from both species. Further, we correct the identification of published genomic/transcriptomic resources that were originally attributed to *O. carmela*, and present new Illumina-sequenced transcriptome assemblies for each of these species, and the mitochondrial genome sequence for *O. pearsei* sp. nov. Using SSU and LSU ribosomal DNA and the mitochondrial genome, we report the phylogenetic relationships of these species relative to other *Oscarella* species, and find strong support for placement of *O. pearsei* sp. nov. in a clade defined by the presence of spherulous cells that contain paracrystalline inclusions; *O. carmela* lacks this cell type and is most closely related to the Western Pacific species, *O. malakhovi. Oscarella pearsei* sp. nov and *O. carmela* can be tentatively distinguished based upon gross morphological differences such as color, surface texture and extent of mucus production, but can be more reliably identified using mitochondrial and nuclear barcode sequencing, ultrastructural characteristics of cells in the mesohyl, and the morphology of the follicle epithelium which surrounds the developing embryo in reproductively active individuals. Usually, cryptic species are very closely related to each other, but in this case and in sponges generally, cryptic species may be very distantly related because sponges can be difficult to identify based upon gross morphological characteristics.

## Introduction

The homoscleromorph sponge species *Oscarella carmela* Muricy & Pearse, 2004 was described from Carmel, California and was the first record of this genus from the Pacific coast of North America [1]. When described, there was question about whether this species was native to the region or whether it was invasive, in part because it was initially observed in public and research aquaria and only later discovered in the nearby intertidal region. Through characterization of its cellular ultrastructure it was ultimately determined to represent a new species, distinct from all known species globally and consistent with the view that it is native to the Eastern Pacific.

We and others were interested in developing this species as a model for genomic and experimental research for several reasons: 1) it is abundant and easily accessible in research aquaria at the Joseph Long Marine Laboratory at the University of California Santa Cruz, 2) in the laboratory environment, embryos of all stages are present year round, albeit more abundant in late summer and fall, and 3) it is thin and therefore internal cells and tissues are easily imaged using common microscopy and experimental methods. To facilitate this development, we sequence expressed sequence tags (ESTs) [2], the mitochondrial genome [3], and a draft nuclear genome [4] for this species.

More recently, we used the Illumina platform to sequence and assemble the transcriptome of *O. carmela* to improve gene prediction from the draft genome, beyond what was possible using ESTs alone. However, from these data (reported in this article) we noticed that there was considerable sequence divergence at both the nucleotide- and amino acid-level between the Illumina transcriptome and previously sequenced ESTs and gene predictions from the draft genome. This led us to suspect that there may be two or more cryptic species that are co-distributed. Importantly, the tissues used to create each dataset were each derived from a single individual, otherwise the presence of multiple cryptic species may have gone undetected.

The original description of *O. carmela* reports the existence of color variants ranging from light brown to orange, and morphological variants ranging from thin and smooth to thicker and having a bumpy, microlobate surface [1]. This is consistent with our personal observations in both the lab and in the field. However, without experience collecting these sponges and without the opportunity to see the various morphotypes side by side, the differences between them appear very subtle and seem to occur along a continuum rather than being discrete (Fig 1). Upon the discovery of such significant differences between different genetic datasets, we were careful to document the morphological differences between individuals and to preserve material for ultrastructural comparison using transmission electron microscopy, and for molecular biology including additional transcriptome sequencing, DNA barcoding and phylogenetic analysis.

**Fig. 1:**
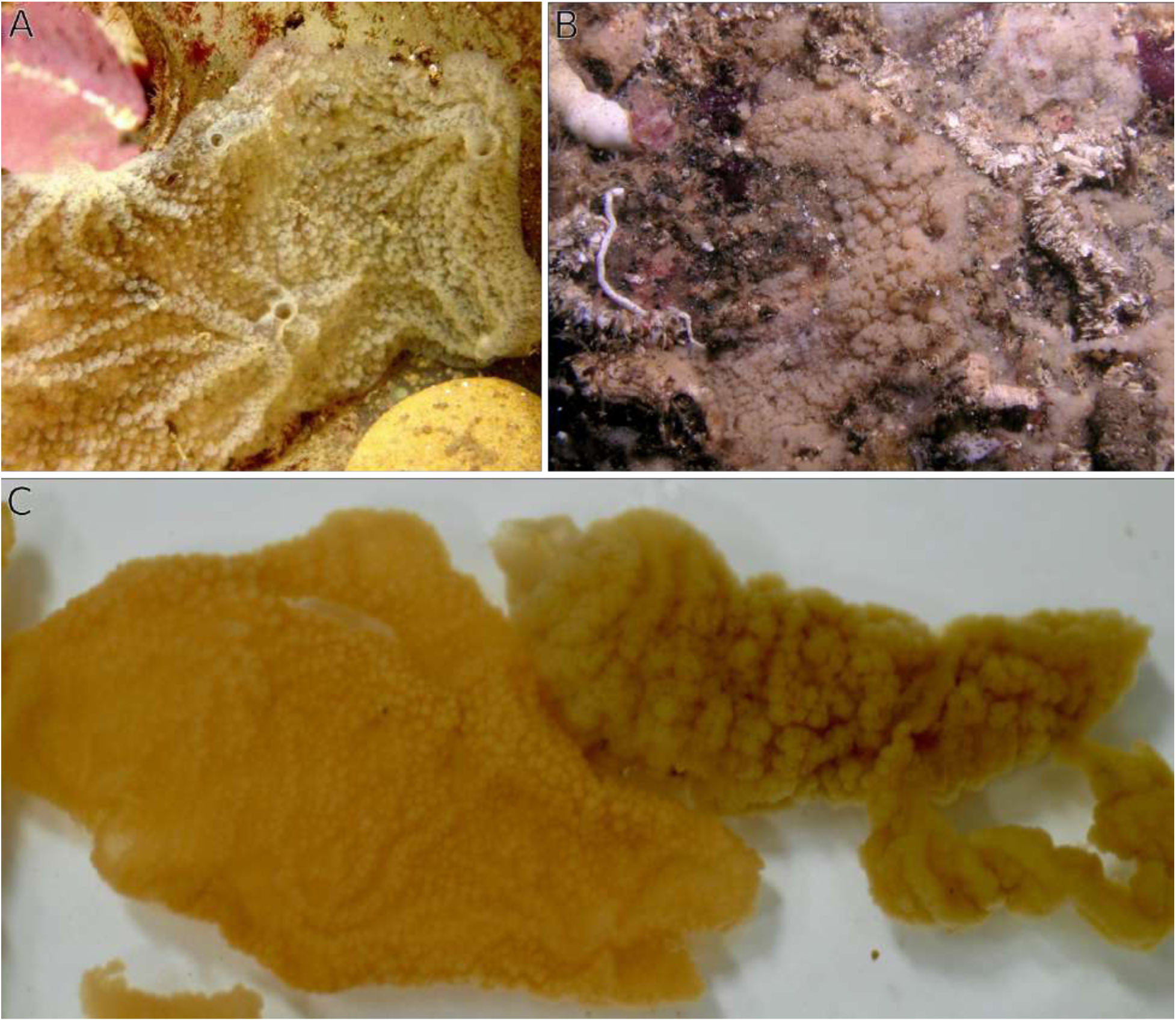
*Oscarella pearsei* sp. nov. and *O. carmela*. **(A)** *In situ* photographs of *O. pearsei* sp. nov. and **(B)** *O. carmela*. **(C)** Upon collection, these species can be distinguished based upon differences in their thickness, surface texture, and color. Left: *O. pearsei* sp. nov., Right: *O. carmela*. (photo credit: J. Mitchell, panel A).

Here we report that there are two species of the genus *Oscarella* co-distributed in the Monterey/Carmel region of central California. Through re-examination of the holotype and paratypes of *O. carmela* we have determined that the original description of this species was based on samples derived from both species. We distinguish between these species, leaving the name *O. carmela* assigned to the species represented by the holotype, and we describe the new species under the name *O. pearsei* sp. nov. In addition to morphological and ultrastructural characterization, we present correctly identified transcriptome assemblies corresponding to each species, their mitochondrial genomes, DNA barcoding sequences that can be used to confirm their identity, and we report their phylogenetic placement relative to each other and other species of *Oscarella*.

## Materials and Methods

### Transcriptome sequencing and Assembly

*Oscarella pearsei* sp. nov. transcriptome assembly (consisting of quality trimming, error correction, *de novo* assembly, identification and removal of cross-contamination, prediction of amino acid sequences, elimination of redundant transcripts, and removal of noise transcripts) was performed according to the protocol described in Peña *et al*., 2016. Within the error correction step, parameter values supplied to the Reptile error correction software v1.1 [5] for the *O. pearsei* sp. nov. read set were as follows (see methods in [6] for a description of how these values were derived): T_expGoodCnt of 54, T_card of 21, KmerLen of 13, Qthreshold of 73, Qlb of 60, MaxBadQPerKmer of 8 and Step to 12, with all other settings at their defaults. The initial *de novo* assembly using Trinity release 2013-02-25 [7] of *O. pearsei* sp. nov. contained 122,656 contigs. The decontamination step removed 385 contigs, with 122,271 contigs remaining. Initial protein prediction on the decontaminated contigs resulted in 146,347 proteins. Redundancy removal based on these predictions eliminated 67,850 redundant contigs and 117,085 proteins, with 54,421 contigs and 29,262 predicted proteins remaining. Subsequent elimination of noise transcripts below FPKM 0.01 resulted in the further removal of 417 contigs containing 42 proteins, resulting in a final *O. pearsei* sp. nov. transcriptome data set of 54,004 contigs and 29,220 predicted proteins.

*Oscarella carmela* transcriptome assembly was performed using essentially the same set of steps, with the following minor modifications (and associated parameter values and contig/protein counts): because *O. carmela* reads were single-end (not paired-end) we performed a single trimming step (equivalent to the second phase of trimming on paired-end data) with Trimmomatic version 0.30 [8], using identical parameter values to *O. pearsei* sp. nov., with the addition of the directive TOPHRED33, as the reads were produced by an earlier generation sequencing machine. During the error correction step, parameter values supplied to Reptile were as follows: T_expGoodCnt of 10, T_card of 4, KmerLen of 13, Qthreshold of 66, Qlb of 58, MaxBadQPerKmer of 8 and Step to 12, with all other settings at their defaults. For initial *de novo* transcriptome assembly with Trinity, we used default parameter values for single-end reads, resulting in 66,109 contigs and 61,705 predicted proteins. We did not perform the removal of cross-contamination step on *O. carmela*, as it was not sequenced on the same flow cell as any other sponges. Redundancy removal eliminated 3,378 contigs and 4,616 proteins, with 62,731 non-redundant contigs and 57,089 non-redundant proteins remaining. We subsequently eliminated 2,893 noise contigs and 3,334 noise proteins below the expression threshold of FPKM of 0.01, resulting in a final *O. carmela* transcriptome data set of 59,838 contigs and 53,755 predicted proteins.

### Mitochondrial genome assembly

Mitochondrial genome of *Oscarella pearsei* sp. nov. was assembled using SOAPdenovo version 1.0 [9] from the Illumina genomic sequences collected in the previous study [4] with a mean coverage around 1500x. The tRNA genes were identified by the tRNAscan-SE program [10]; rRNA and protein genes were identified by similarity searches in local databases using the FASTA program [11]. The presence of inversion was confirmed by mapping individual Illumina sequences on mitochondrial genome assembly and by conducting PCR amplifications across the inversion boundaries.

### Phylogenetic Analyses

Phylogenetic analyses of SSU and LSU data were performed on the phylogeny.fr platform [12] and comprised the following steps. Sequences were aligned using CLC Workbench (v. 6.8) (Qiagen). After alignment, ambiguous regions (i.e., containing gaps and/or poorly aligned) were removed with Gblocks (v0.91b; [13]) using the following parameters: -minimum length of a block after gap cleaning: 10, -no gap positions were allowed in the final alignment, - all segments with contiguous non-conserved positions bigger than 8 were rejected, -minimum number of sequences for a flank position: 85%. Phylogenetic trees were reconstructed using the maximum likelihood method implemented in the PhyML program (v3.1/3.0 aLRT) [14]. The default substitution model was selected assuming an estimated proportion of invariant sites (of 0.357) and 4 gamma-distributed rate categories to account for rate heterogeneity across sites. The gamma shape parameter was estimated directly from the data (gamma=0.635). Reliability for internal branches was assessed using the aLRT test (SH-Like) [15]. Graphical representation and edition of the phylogenetic trees were performed with TreeDyn (v198.3) [16].

Inferred amino acid sequences of individual mitochondrial proteins from homoscleromorph sponges were aligned with Mafft v.6.861b [17]. Conserved blocks within the alignments were selected with Gblocks 0.91 b [13] using relaxed parameters (parameters 1 and 2 = ½, parameter 3 = 8, parameter 4 = 5, all gap positions in parameter 5). Cleaned alignments were concatenated in a dataset 4070 positions in length. Bayesian inferences based on amino acid sequences were performed with the CAT+GTR+G4 mixture model implemented in the program PhyloBayes MPI version 1.4e [18]. PhyloBayes analysis consisted of four chains over 57,000 generations (max diff. ∼0.03). The chains were sampled every tenth generation after the first 100 burn-in cycles.

### DNA barcoding

Upon collection, an ∼1cm x 1cm piece of tissue from each individual was preserved in fixative for Transmission Electron Microscopy (described below) and in 95% EtOH for genomic DNA extraction and DNA barcoding. Genomic DNA for LSU barcoding was isolated using the GenElute Mammalian Genomic DNA Miniprep kit (Sigma-Aldrich) per manufacturer specifications (EtOH was allowed to evaporate before starting procedure). Using 28S-C2-fwd and 28S-D2-rev [19] primers, a fragment of the LSU was amplified by PCR and sequenced by Eurofins Genomics (Germany). Genomic DNA for mitochondrial barcoding was isolated using the standard phenol-chloroform method [20]. Using diplo-cob-f1m and diplo-cob-r1m primers [21], a fragment of *cob* was amplified by PCR (Invitrogen recombinant *Taq* DNA polymerase kit), purified with Promega Wizrd SV Gel and PCR Clean-up system, and sequenced by Iowa State University Sequencing facility.

### Transmission Electron Microscopy

For semi-thin sections and for transmission electron microscopy (TEM) investigations, sponges were fixed in a solution composed of one volume of 25% glutaraldehyde, four volumes of 0.2 M cacodylate buffer and five volumes of filtered seawater for at least 2 h before being post-fixed in 2% OsO_4_ in seawater at room temperature for 2 h. After fixation, samples were washed in 0.2 M cacodylate buffer and distilled water successively, and dehydrated through a graded ethanol series. Specimens were embedded in Araldite resin. Semi-thin sections (1 μm in thickness) were cut on a Reichert Jung ultramicrotome equipped with a “Micro Star” 45° diamond knife before being stained with toluidine blue, and observed under a WILD M20 microscope. Digital photos were taken with a Leica DMLB microscope using the Evolution LC color photo capture system. Ultrathin sections (60 – 80 nm) were cut with a Leica UCT ultramicrotome equipped with a Drukkert 45° diamond knife. Ultrathin sections, contrasted with uranyl acetate, were observed under a Zeiss-1000 transmission electron microscope (TEM).

## Results

The holotype and paratypes submitted as part of the original species description for *Oscarella carmela* we re-examined and found to include tissue from distinctly separate species (Table 1) based upon distinguishing characteristics defined below. The decision as to which species should retain the name *O. carmela* was made based upon which corresponded to the holotype submitted to the Museu Nacional Rio de Janeiro (MNRJ), and the other species was described as *O. pearsei* sp. nov. Thus, previously published studies of the species *O. carmela* may actually refer to either of these co-distributed species. In Table 2 we provide a list of previously published and new genomic and transcriptomic resources, and clarify the species from which they were derived.

**Table 1.** Original *O. carmela* holotype and paratypes submitted to MNRJ.

| Accession number | Re-examined and identified as: |
| --- | --- |
| Holotype B-6618 #2 | *O. carmela* |
| Paratype B-6617 #1 | *O. carmela* |
| Paratype B-6616 | *O. pearsei* sp. nov. |

**Table 2.** Material source for genomic and transcriptomic data.

| Description | Corrected Species Designation | Reference |
| --- | --- | --- |
| ESTs from mixed adult and embryonic tissues (Illumina) | *O. pearsei* sp. nov. | Nichols et al. 2006 |
| Draft genome assembly (Illumina) | *O. pearsei* sp. nov. | Nichols et al. 2012 |
| mtGenome | *O. carmela* | Wang et al. 2007 |
| mtGenome | *O. pearsei* sp. nov. | This study |
| Transcriptome from adult tissues (Illumina) | *O. carmela* | This study |
| Transcriptome from adult tissues (Illumina) | *O. pearsei* sp. nov. | This study |

### Transcriptome sequencing

Transcriptome sequencing was performed on adult somatic tissues from a single individual of the species *O. pearsei* sp. nov., and on tissues and brooded embryos derived from a single individual of *O. carmela*. Assembly of the *O. carmela* transcriptome resulted in 59,838 contigs encoding 53,755 predicted peptides of >50aa in length. Assembly of the *O. pearsei* sp. nov. transcriptome resulted in 54,004 contigs encoding 29,220 predicted peptides of >50aa in length. The difference in number of peptides predicted from these datasets was explored using CD-HIT [22] to create peptide clusters with high sequence identity. *O. carmela* peptides formed 37,887 unique clusters whereas *O. pearsei* sp. nov. peptides formed 29,209 unique clusters. These results are consistent with the fact that the *O. carmela* transcriptome was assembled from fewer reads than the *O. pearsei* sp. nov. assembly, and differences in transcript number may result from assembly errors due to low sequencing coverage that mimic alternative splice variants. Raw sequence data are available at the NCBI Sequence Read Archive under the accession numbers SRX388205 and SRX386257. Assembled transcriptomes and predicted peptides are available at compagen.org [23].

### Mitochondrial genome

The mitochondrial genome for *O. carmela* was previously published [3], and the species designation for this dataset remains unchanged. Here, we report and describe the sequence of the mitochondrial genome for *O. pearsei* sp. nov. (Fig 2), which was assembled from Illumina DNAseq data and deposited to GenBank (accession number KY682864). The mitochondrial genome of *O. pearsei* sp. nov. is a circular mapping molecule 20,320 bp in size and 65.6% A+T that fits well with the description of a typical mitochondrial genomes in the family Oscarellidae [24, 25]. It contains 42 genes, organized in two clusters with opposite transcriptional polarities. The genes include the unusual *tatC*, for subunit C of the twin arginine translocase [26] as well as a duplicated *trnV*(*uac*). However, mitochondrial gene order in *O. pearsei* sp. nov. is unique among other representatives of the family Oscarellidae due to an inversion of a mtDNA fragment between the two copies of the *trnV* gene. This inversion allows an easy PCR-based identification of two sympatric species of *Oscarella*. In addition, the second duplicated tRNA gene (most likely *trnT*) encodes an unusual tRNA-like structure with well conserved aminoacyl acceptor, TYC and DHU arms, but with the anticodon arm replaced by a 9-nt loop. An identical structure with a very similar sequence (two substitutions) is also encoded in the mtDNA of the closely related *O. balibaloi* (accession number KY682865). Interestingly, in the latter species the copy of trnV between *atp8* and *nad1* encodes a similar structure with identically reduced anticodon arm. The mt-coding sequences of *O. carmela* and *O. pearsei* sp. nov. are on average 89.4% identical with the rate of synonymous substitutions estimated between 0.31 and 2.75 (mean=0.76) and the rate of non-synonymous substitutions estimated between 0.01 and 0.21 (mean=0.04). Both synonymous and non-synonymous substitution rates were highest in *tatC*, an observation consistent with our previous study [26].

**Fig. 2.**
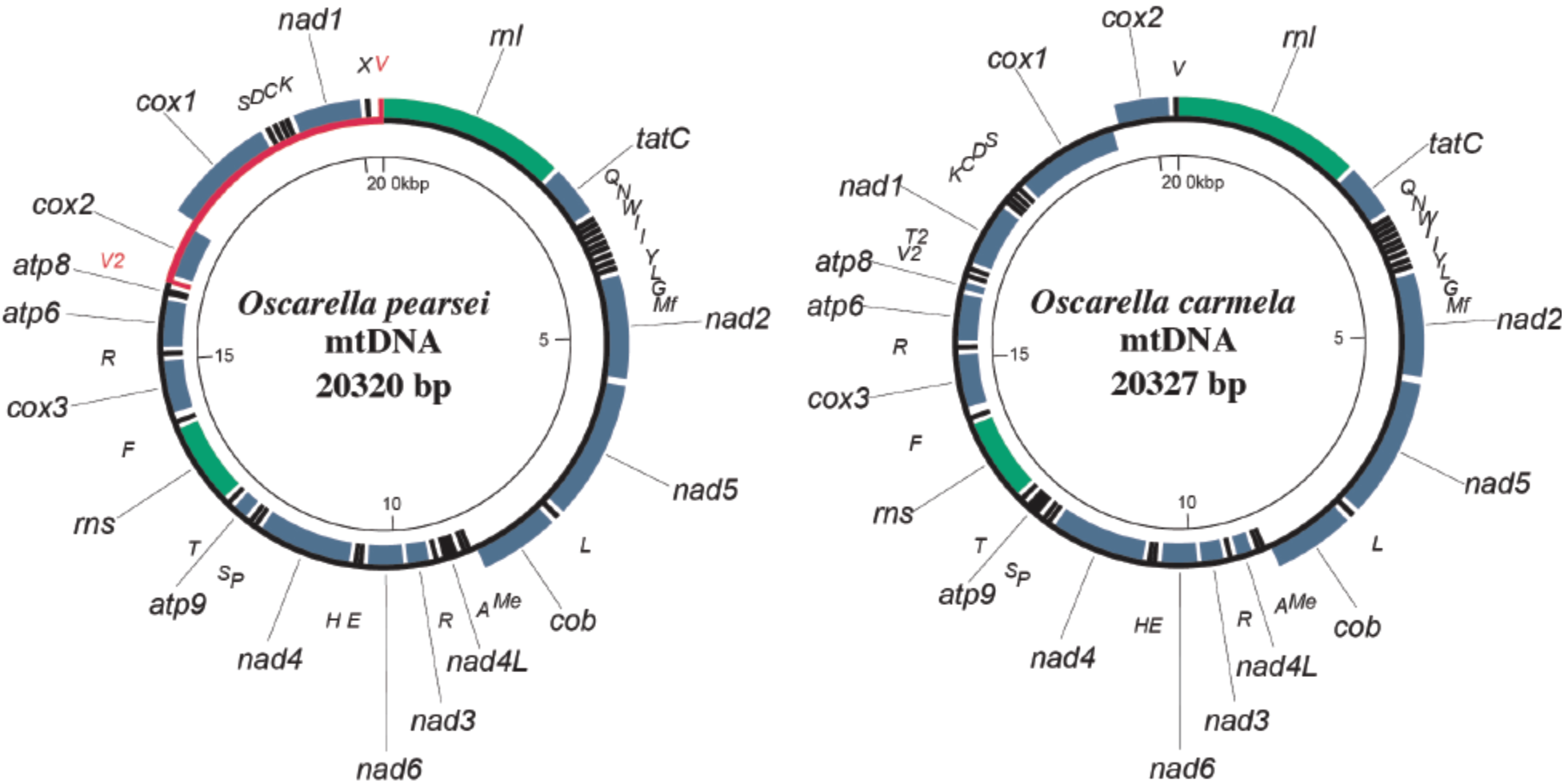
Phylogenetic analysis using ribosomal DNA data. Phylogenetic trees constructed using SSU/LSU rDNA and mitochondrial genome sequences from *Oscarella* and outgroup species.

### Phylogenetic analysis

To assess the relationship of *O. carmela* and *O. pearsei* sp. nov., to each other and to other known *Oscarella* species globally, phylogenetic analyses were performed using three datasets: small subunit (SSU), large subunit (LSU) rDNA and the whole mitochondrial genome. Maximum likelihood and Bayesian analyses of each dataset were highly concordant and collectively supported the close relationship of *O. carmela* to *O. malakhovi*, and the placement of *O. pearsei* sp. nov. in a subclade containing *O. kamchatkensis, O. nicolae, O. balibaloi*, and *Pseudocorticium jarrei* (Fig 3).

**Fig. 3.**
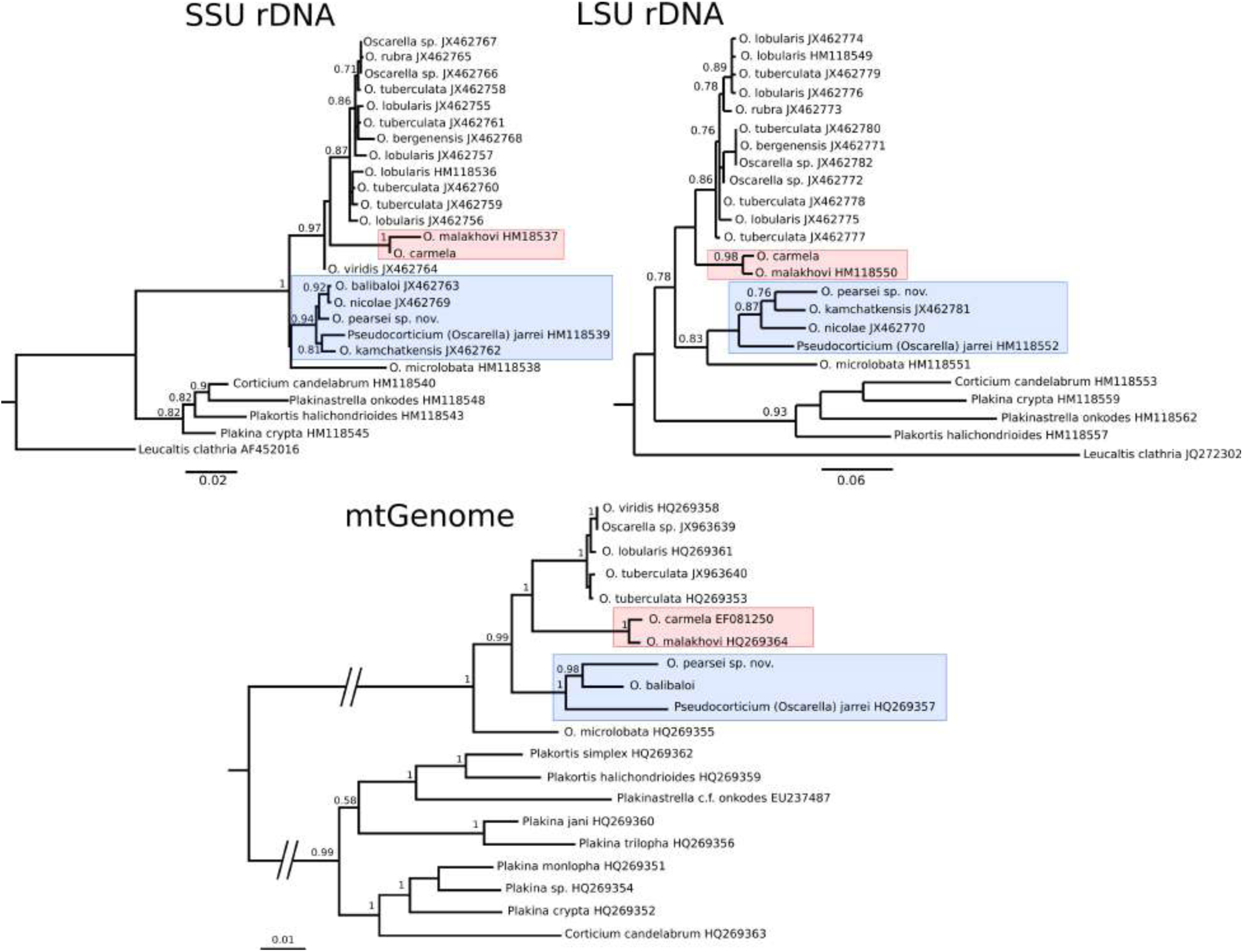
Mitochondrial Genome Architecture. Mitochondrial genome maps of *Oscarella pearsei* sp. nov. and *O. carmela*. Protein (green) and ribosomal (blue) genes are *atp6, atp8-9*: subunites 6, 8, and 9 of F0 adenosine triphosphatase (ATP) synthase; *cob*: apocytochrome b; *cox 1-3*: cytochrome c oxidase subunites 1-3; *nad1-6* and *nad4L*: NADH dehydrogenase subunits 1-6 and 4L; *rns* and *rnl*: small and large subunit rRNAs; *tatC*: twin-arginine translocase component C. tRNA genes (black) are identified by the one-letter code for their corresponding amino acid. Genes outside the main circle are transcribed clock-wise, inside - counter clock-wise. Large inversion between two copies of *trnV* in *O. pearsei* sp. nov. mtDNA is shown in red.

### DNA barcoding

All museum-deposited samples used for the re-description of *O. carmela* and the new description of *O. pearsei* sp. nov. were identified using a short, rapidly evolving region of the LSU that has been shown to be useful for DNA barcoding [19, 27] as well as a fragment of mitochondrial Cytochrome b gene [21]. Both LSU and *cob* fragments were found to reliably distinguish between the two species, which were detected living side by side in the laboratory (UCSC and at the type locality in the intertidal zone of Carmel, CA). The *O. carmela* LSU barcode sequence was deposited to GenBank under accession KY513287 and the *O. pearsei* sp. nov. barcode sequence was deposited to GenBank under accession KY513286. The barcode sequence is 458bp in *O. pearsei* sp. nov. and 446bp in *O. carmela. Cob* barcoding sequences were found to be identical to the fragments of *cob* in complete mitochondrial genomes of the two species. No polymorphisms were detected between individuals sampled from within each species (n=5) in either nuclear or mitochondrial barcodes and the sequence identity for these markers between the two species was 77% and 87%, respectively.

### Morphological descriptions

Genus *Oscarella* Vosmaer, 1884

#### Type species

*Halisarca lobularis* Schmidt, 1862 (by monotypy). [*Oscaria*] Vosmaer, 1881: 163 (preocc. by *Oscaria* Gray, 1873 – Reptilia); *Oscarella* Vosmaer, 1884: pl. 8 (explanation); 1887: 326 (nom. nov. for *Oscaria* Vosmaer). *Octavella* Tuzet and Paris, 1964: 88.

#### Diagnosis (modified from Muricy & Diaz 2002)

Homoscleromorpha without skeleton, with a variable degree of ectosome development. The aquiferous system has a sylleibid-like or leuconoid organization, with eurypylous or diplodal choanocyte chambers. Mitochondrial genomes that have been sequenced so far encode a gene absent in other animal mitochondrial genomes: *tatC*.

#### *Oscarella pearsei* sp. nov

urn:lsid:zoobank.org:act:8431B41C-B08E-4205-8FE6-CCCFFB6EB574

#### Holotype

ZIN RAS №№ 11798, Ethanol. urn:lsid:zoobank.org:act:8431B41C-B08E-4205-8FE6-CCCFFB6EB574, sea tables at Long Marine Lab. Santa Cruz, California, USA, Collected by Scott Nichols 01.08.2015.

ZIN RAS № 11800, Glutaraldehyde. urn:lsid:zoobank.org:act:8431B41C-B08E-4205-8FE6-CCCFFB6EB574, sea tables at Long Marine Lab. Santa Cruz, California, USA, Collected by Scott Nichols 01.08.2015.

#### Paratypes

ZIN RAS № 11799, Ethanol, sea tables at Long Marine Lab. Santa Cruz, California, USA, Collected by Scott Nichols 01.08.2015.

ZIN RAS № 11801, Glutaraldehyde, sea tables at Long Marine Lab. Santa Cruz, California, USA, Collected by Scott Nichols 01.08.2015.

RBINS-IG 33439-POR.0122, Ethanol, sea tables at Long Marine Lab. Santa Cruz, California, USA, Collected by Scott Nichols 01.08.2015.

RBINS-IG 33439-POR.0123, Glutaraldehyde, sea tables at Long Marine Lab. Santa Cruz, California, USA, Collected by Scott Nichols 01.08.2015.

RBINS-IG 33439-POR.0124 Glutaraldehyde, Ethanol, sea tables at Long Marine Lab. Santa Cruz, California, USA, Collected by Scott Nichols 01.08.2015.

#### Comparative material examlned

*Oscarella carmela*, Holotype MNRJ 8033 (slide B-6618 #2), Carmel Point, Carmel Bay, intertidal, Collected by J. Pearse, 08. 05. 2003 [1]. *Oscarella carmela* Paratype, CASIZ 168925 (slide B-6617 #1), Carmel Point, Carmel Bay, central California, intertidal, Collected by J. Pearse. 08. 05. 2003 [1]. *Oscarella carmela* Muricy, Pearse, 2004. Syntype: ZIN RAS № 11802, Ethanol (this study), sea tables at Long Marine Lab. Santa Cruz, California, USA, S. Collected by Scott Nichols 01.08.2015. *Oscarella carmela* Muricy, Pearse, 2004. Syntype, ZIN RAS № 11803, Ethanol (this study), sea tables at Long Marine Lab. Santa Cruz, California, USA, S. Collected by Scott Nichols 01.08.2015. *Oscarella carmela* Muricy, Pearse, 2004. Syntype ZIN RAS № 11804, Glutaraldehyde (this study), sea tables at Long Marine Lab. Santa Cruz, California, USA, S. Collected by Scott Nichols 01.08.2015. *Oscarella carmela* Muricy, Pearse, 2004. Syntype ZIN RAS № 11805, Glutaraldehyde (this study), sea tables at Long Marine Lab. Santa Cruz, California, USA, S. Collected by Scott Nichols 01.08.2015.*Oscarella carmela* Muricy, Pearse, 2004. Syntype RBINS-IG 33439-POR.0125, Ethanol (this study), sea tables at Long Marine Lab. Santa Cruz, California, USA, S. Collected by Scott Nichols 01.08.2015. *Oscarella carmela* Muricy, Pearse, 2004. Syntype RBINS-IG 33439-POR.0126, Ethanol (this study), sea tables at Long Marine Lab. Santa Cruz, California, USA, S. Collected by Scott Nichols 01.08.2015. *Oscarella carmela* Muricy, Pearse, 2004. Syntype RBINS-IG 33439-POR. 0127, Glutaraldehyde (this study), sea tables at Long Marine Lab. Santa Cruz, California, USA, S. Collected by Scott Nichols 01.08.2015. *Oscarella carmela* Muricy, Pearse, 2004. Syntype, RBINS-IG 33439-POR.0128, Glutaraldehyde (this study), sea tables at Long Marine Lab. Santa Cruz, California, USA, S. Collected by Scott Nichols 01.08.2015. *Oscarella malakhovi*

Ereskovsky, 2006 (ZIN RAS 10697 ZIN RAS 10698: Japan Sea) [28]. *Oscarella kamchatkensis* Ereskovsky, Sanamyan & Vishnyakov, 2009 (ZIN RAS 11058, ZIN RAS 11059 and ZIN RAS 11060: North Pacific, Avacha Gulf) [29]. *Oscarella lobularis* (Schmidt, 1862) and *Oscarella tuberculata* (Schmidt, 1868). NE Mediterranean Sea (Marseille region), underwater cave of Maire Island. *Oscarella microlobata* Muricy, Boury-Esnault, Bézac, Vacelet, 1996 and *Oscarella viridis* Muricy, Boury-Esnault, Bézac, Vacelet, 1996. NE Mediterranean Sea (Marseille region), underwater cave of Jarre Island [30]. *Oscarella balibaloi*, Pérez, Ivanišević, Dubois, Pedel, Thomas, Tokina, Ereskovsky, 2011. NE Mediterranean Sea (Marseille region), underwater cave of Maire Island [31]. *Oscarella nicolae* Gazave, Lavrov, Cabrol, Renard, Rocher, Vacelet, Adamska, Borchiellini, Ereskovsky, 2013 (MNHN DJV155, LSID urn:lsid:zoobank.org:act:DFFAD94B-9CAD-4F99-994D-BDEF75EF98A1, MNHN DJV156) North Sea, Norway (Bergen Fjords, Skarvoysundet) [25]. *Oscarella bergenensis* Gazave, Lavrov, Cabrol, Renard, Rocher, Vacelet, Adamska, Borchiellini, Ereskovsky, 2013 (MNHN DJV153, LSID:urn:lsid:zoobank.org:act:2D44BCFA-2163-47C7-9E70-EF6C13E0E4A4, MNHN DJV154) North Sea, Norway (Bergen Fjords, Skarvoysundet) [25].

#### Diagnosis

Found on the sides and undersides of rocks in the intertidal zone in Carmel, California, USA. Encrusting *Oscarella*, tan to orange in color, soft, slimy but delicate consistency with two particular mesohylar cell types with inclusions: abundant granular cells and spherulous cells with paracrystalline inclusions. Nucleus in choanocyte has apical position. Embryo follicle layer comprised of multilayer cubic cells. It has only one morphotype of endobiotic bacteria.

#### Description

Sponge habit is very thinly encrusting. Size can vary, with most sponges being less than 10 cm in diameter, but sometimes extending to 30 cm or greater. Color *in vivo* is light tan to orange (Fig 1A). Preserved fragments are lighter in color, sometimes approaching white. The surface is finely and uniformly lobate with small hemispherical lobes, and adorned with few delicate oscula 3-5 mm in height; oscula are only visible in submerged specimens that are relaxed (i.e., not recently contracted). Sponges are not uniformly attached to the substrate; rather, outgrowths extend from the basopinacoderm and attach to the substrate at their tip. Thus, the sponge can be easily dislodged from the substrate and maintains its integrity.

#### Soft tissue organization

Spicule and fiber skeleton absent. Ectosome is between 8 to 15 μm thick. Inhalant canals run perpendicular to the surface. Choanocyte chambers ovoid to spherical, eurypilous, from 55 to 70 μm in diameter. Exhalant canals run towards a well developed system of basal cavities leading to the oscula.

### Cytology

**Exopinacocytes** (Fig 4A) flagellated with flat to oval shape, 13.5 μm wide by 4.9 μm high (N = 10). Cytoplasm containing electron dense granules from 0.6 to 1.2 μm in diameter and electron translucent vacuoles 0.2 – 0.7 μm in diameter. Nucleus about 2.8 μm in diameter with central position. External surface of exopinacocytes covered by a layer of glycocalyx 0.15 - 0.25 μm thick and basal surface has filopodia that extend into the extracellular matrix (S1 Figure).

**Fig. 4.**
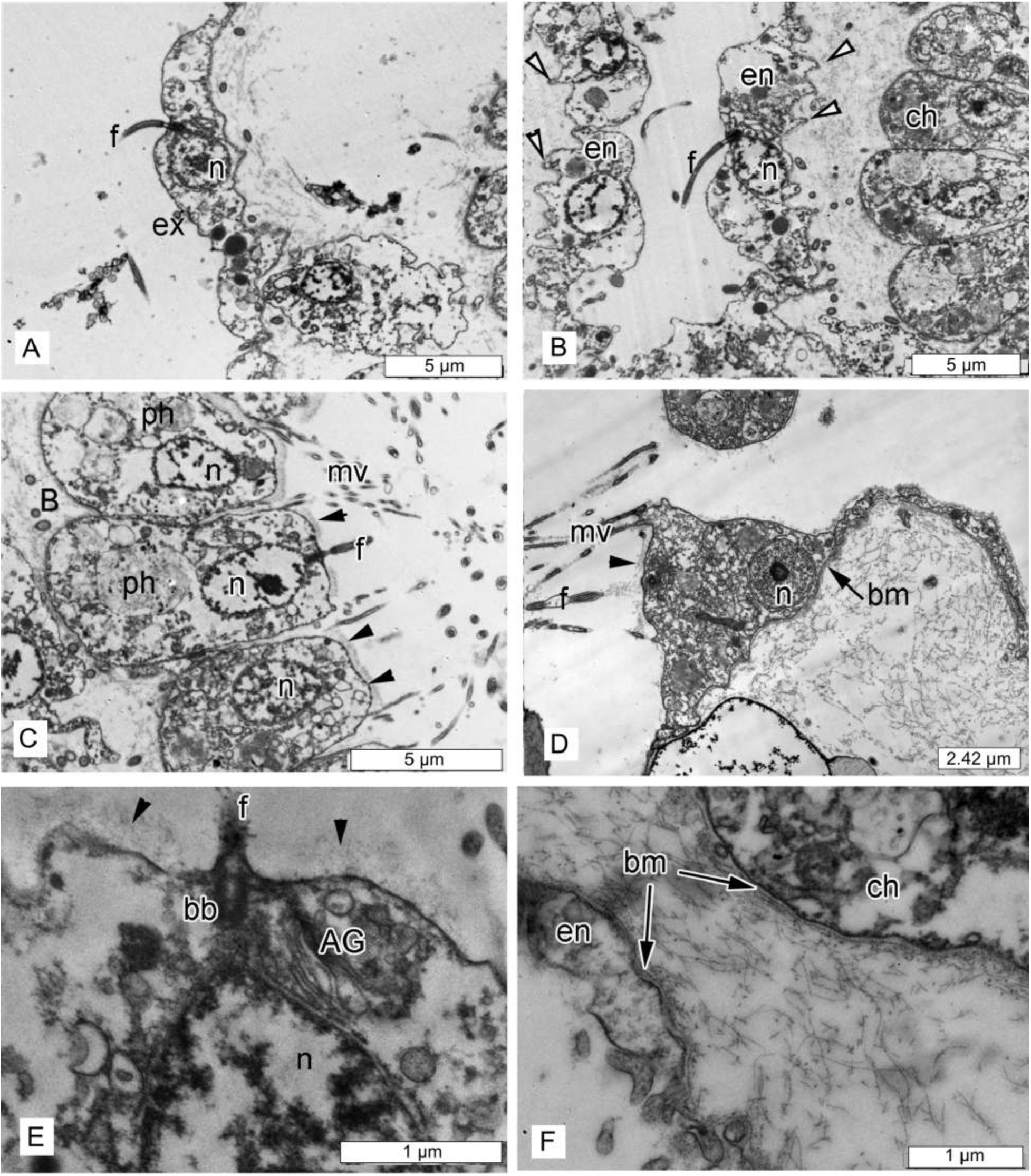
*Oscarella pearsei* sp. nov. TEM of cells and symbiotic bacteria. **(A)** exopinacocyte; **(B)** endopinacocyte; **(C)** choanocytes; **(D)** apopylar cell; **(E)** apical part of choanocyte with glycocalyx layer; **(F)** basal parts of choanocyte and endopinacocyte with basement membrane. (Abbreviations: AG – Golgi apparatus, ap – apopylar cell, B – symbiotic bacteria, bb – basal body of flagellum, bm – basement membrane, ch – choanocytes, en – endopinacocyte, ex – exopinacocyte, f – flagellum, mv – microvilli, n – nucleus, ph – phagosome. Arrowheads – white: basal surface has pseudopodia that extend into the extracellular matrix. black: glycocalyx).

**Endopinacocytes** (Fig 4B) flagellated flat cells, about 17.2 μm wide by 3.3 μm high (N = 10). Nucleus ovoid (2.6 μm in diameter). Cytoplasm includes electron dense granules 0.3 - 0.8 μm in diameter and electron translucent vacuoles 0.3 - 0.8 μm in diameter. Basal surface has pseudopodia that extend into the extracellular matrix.

**Choanocytes** (Fig 4B, C and E) ovoid to trapezoidal, about 4.2 μm wide by 7.7 μm high (N = 20). Nucleus (∼2.4μm in diameter) has apical position and contacts with flagellar basal apparatus. Cytoplasm includes from one to eight phagosomes 1.2 – 3.4 μm in diameter.

**Apopylar cells** (Fig 4D) roughly triangular or oval in section, 6.3 μm wide by 4.7 μm high (N = 5). Nucleus up to 2.9 μm in diameter. Cytoplasm contains mitochondria, digestive vacuoles, and small osmiophilic inclusions.

Surface of endopinacocytes, choanocytes and apopylar cells covered by thin, irregular glycocalyx, choanoderm and pinacoderm underlined by continuous basement membrane (Fig 4D-F).

#### Two types of cells with inclusions occur within the mesohyl

**Granular cells** (Fig 5A) ovoid to elongate, approximately 5.4 × 7.1 μm, sometimes amoeboid-like or irregular (N = 8). Cytoplasm filled with oval granules. Some granular cells exclusively contain osmiophilic granules that have a homogenous, opaque (gray) appearance, whereas others have fewer of this variety (or none) in combination with vacuoles (1.3 - 2.8 μm in diameter) that contain filamentous content. This seems to reflect a dynamic process of vacuolar content maturation. At the very least, intermediate phenotypes are apparent, thus we presume that these represent a single type of granular cell (S2 Fig). Nucleus regular, about 2.2 μm in diameter without nucleolus. During reproduction, these cells concentrate around, and incorporate into the eggs after closing of the follicle (S3 Fig).

**Fig. 5.**
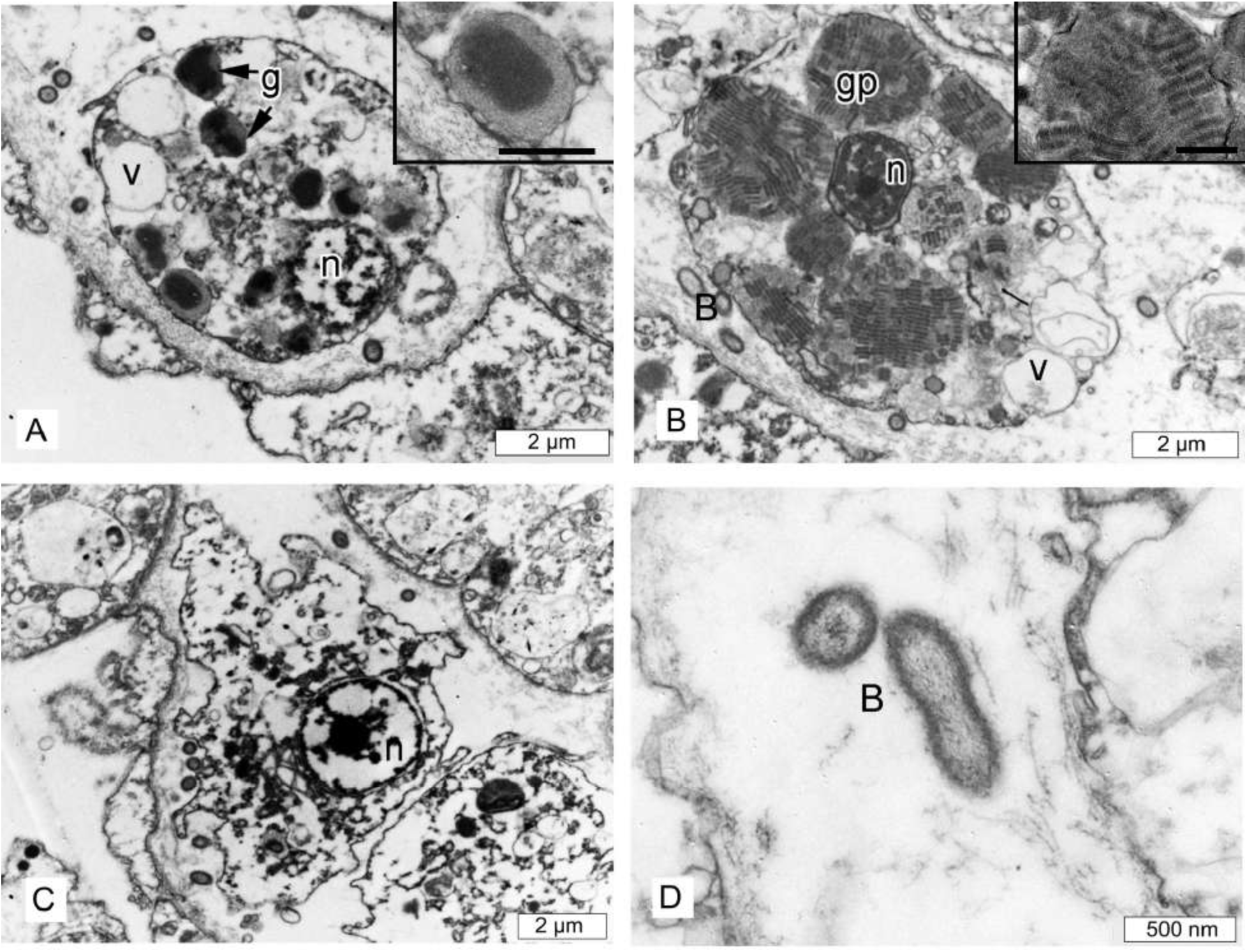
*Oscarella pearsei* sp. nov. mesohylar cells. **(A)** granular cell type 1; inset - detail of granule, scale bar = 0.5 μm; **(B)** spherulous cells with paracrystalline inclusions; inset - detail of granule, scale bar = 0.5 μm; **(C)** archeocyte; **(D)** symbiotic bacteria. (Abbreviations: B – symbiotic bacteria, g – granule, gp - granule with paracrystalline inclusions, n – nucleus, v – vacuole).

**Spherulous cells with paracrystalline inclusions** (Fig 5B). Ovoid or spherical cells 9.2 μm long and 8.2 μm in diameter with nucleolated nucleus 2.2 μm in diameter. Cytoplasm is filled with 6 - 11 spherical heterogeneous inclusions in section (1.9 - 2.8 μm in diameter), composed of paracrystalline elements included in a homogenous matrix. Paracrystalline elements are ovoid or cylindrical in longitudinal section and round in transversal section (0.7 μm long and 0.23 μm in diameter). These elements are composed of fibrils arranged in a transverse banding pattern with dark bands alternated by clear bands. In cross sections, the paracrystalline elements are organized in spiral lines. Cytoplasm can also contain electron transparent vacuoles (0.9 – 1.6 μm in diameter). These cells also include vacuoles with irregular shape and homogenous, opaque (gray) contents from 0.3 to 1.1 μm, internal periphery of these vacuoles have electron dense thickening. During reproduction, these cells concentrate around the eggs and are located inside them after the closing of follicle (S3 Fig).

**Archeocytes** (Fig 5C) are amoeboid, 7.3 μm wide by 5.0 μm long, and dispersed in low number in the mesohyl. Their cytoplasm includes small osmiophilic inclusions from 0.2 to 0.6 μm in diameter. The nucleus is spherical or ovoid and ∼2.4 μm in diameter and often include a nucleolus.

**Endobiotic bacteria** (Fig 5D) One morphotype of extracellular endobiotic bacteria occurs in the mesohyl: B1. These bacteria are abundant, rod-like, slightly curved shape 0.74 – *0,85* - 1.0 μm in length and 0.34 - *0.35* - 0.37 μm in diameter (N = 22). The cell wall consists of two membranes, and a tight periplasmic space. The cytoplasm is filled with filamentous materials between the cell wall and nucleoid. Nucleoid filamentous network is irregular, with thick elements in the center and thin filaments closer to the periphery. Surface covered with thin filamentous outgrowths. These endobiotic bacteria are not abundant.

#### Reproduction

*Oscarella pearsei* sp. nov. is a viviparous and hermaphroditic species; male and female reproductive elements are present in the same individuals. The spermatic cysts rare within the mesohyl of all investigated sponges. They are oval to spherical in shape, with a diameter of about 35 μm and are randomly distributed in the sponge choanosome (Fig 6A). Spermatogenesis is generally asynchronous inside spermatic cysts.

**Fig. 6.**
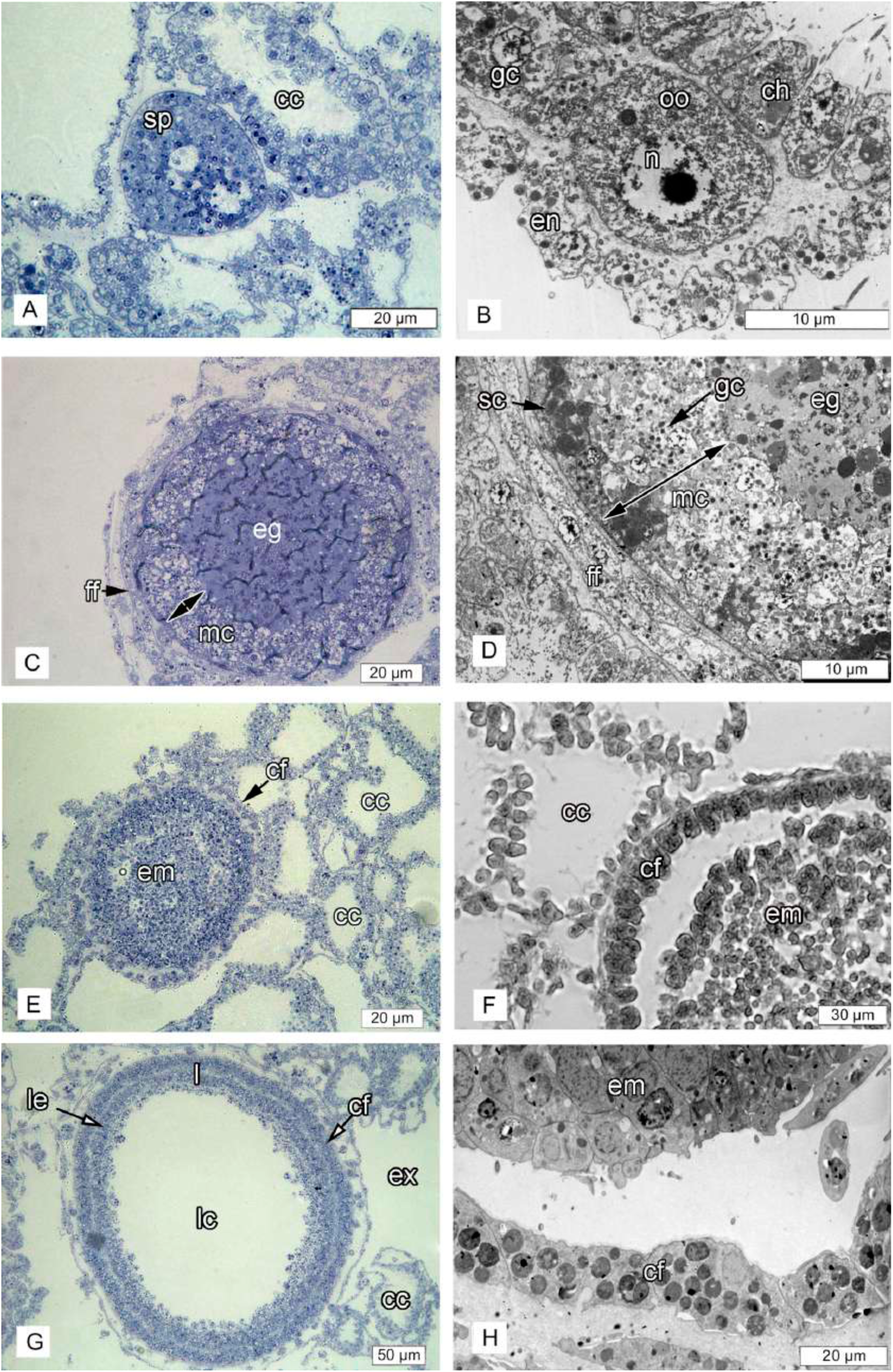
*Oscarella pearsei* sp. nov. reproduction. **(A)** semi-thin section of the spermatocyst; **(B)** TEM of young oocyte before vitellogenesis; **(C)** semi-thin section of the egg with flat monolayer follicle and the thick layer of maternal cells; **(D)** TEM of flat monolayer follicle of the egg and the thick layer of maternal cells; **(E)** semithin section of the embryo with bilayer follicle composed of cubical cells; **(F)** histological section of embryos with cubical follicle; **(G)** semi-thin section of the cinctoblastula larva; **(H)** TEM of the embryo with bilayer follicle with flat external layer and cubical internal ones. (Abbreviations: cc - choanocyte chamber, cf – cubic follicle, ch – choanocyte, eg – egg, em – embryo, en – endopinacocyte, ex – exhalant canal, ff – flat follicle, gc – granular cells; l – larva, lc – larval cavity, le – larval ciliated epithelium, mc – maternal cells; n – nucleus, oo-oocyte, sc – spherulous cells with paracrystalline inclusions, sp – spermatocyst).

Oogenesis and embryogenesis are asynchronous: all stages from oogonia to egg were observed within the same sponge. Young oocytes have an ovoid or amoeboid shape and are situated between choanocyte chambers and endopinacoderm (Fig 6B). Mature eggs are about 70 μm in diameter, isolecithal and polylecithal, with a cytoplasm full of yolk granules (Fig 6C). Embryogenesis is also asynchronous leading to formation of typical cinctoblastula larvae (Fig 6E and G). All stages from cleaving embryos to larva were observed from mid-June to September. Eggs and embryos are located in the basal part of the choanosome and are completely surrounded by a follicle. From the stage of vitellogenesis of the oocyte to the embryos/larva stage the follicle transforms from flat monolayer composed of pinacocytes (oogenesis) (Fig 6C and D) to cubical one in embryos and larva (Fig 6E-H). At the late stages of vitellogenesis the area between the follicle and oocyte contains a thick layer (about 20 – 25 μm) of maternal cells (granular and spherulous cells with paracrystalline inclusions) (Fig 6C and D; S3 Fig). Over the course of embryonic development this layer disappears (S3 Fig).

#### Habitat

*Oscarella pearsei* sp. nov. is found in Carmel, California (36° 48’ N 121° 54’ W) in the low intertidal zone, on the underside of rocks. This is the only location where it has been collected in the field, and is found growing alongside *Oscarella carmela*. Otherwise, both species are found growing in public aquaria and at research facilities along the central California coast (UCSC Long Marine Lab and the Monterey Bay Aquarium) that maintain flow-through seawater tanks that are supplied with seawater pumped directly from the ocean.

#### Etymology

The specific name is given in honor of University of California Santa Cruz Professor Emeritus John Pearse for his support while describing this sponge.

### Redescription of *Oscarella carmela* Muricy, Pearse, 2004

#### Type material

##### Holotype

MNRJ 8033 (slide B-6618 #2), Carmel Point, Carmel Bay, intertidal, Collected by J. Pearse, 08. 05. 2003 [1].

##### Paratypes

CASIZ 168925 (slide B-6617 #1), Carmel Point, Carmel Bay, central California, intertidal, Collected by J. Pearse. 08. 05. 2003 [1].

##### Syntypes

*Oscarella carmela* Muricy, Pearse, 2004. ZIN RAS № 11802, Ethanol (this study), sea tables at Long Marine Lab. Santa Cruz, California, USA, S. Collected by Scott Nichols 01.08.2015. *Oscarella carmela* Muricy, Pearse, 2004. ZIN RAS № 11803, Ethanol (this study), sea tables at Long Marine Lab. Santa Cruz, California, USA, S. Collected by Scott Nichols 01.08.2015. *Oscarella carmela* Muricy, Pearse, 2004. ZIN RAS № 11804, Glutaraldehyde (this study), sea tables at Long Marine Lab. Santa Cruz, California, USA, S. Collected by Scott Nichols 01.08.2015. *Oscarella carmela* Muricy, Pearse, 2004. ZIN RAS № 11805, Glutaraldehyde (this study), sea tables at Long Marine Lab. Santa Cruz, California, USA, S.

Collected by Scott Nichols 01.08.2015. *Oscarella carmela* Muricy, Pearse, 2004. RBINS-IG 33439-POR.0125, Ethanol (this study), sea tables at Long Marine Lab. Santa Cruz, California, USA, S. Collected by Scott Nichols 01.08.2015. *Oscarella carmela* Muricy, Pearse, 2004. RBINS-IG 33439-POR.0126, Ethanol (this study), sea tables at Long Marine Lab. Santa Cruz, California, USA, S. Collected by Scott Nichols 01.08.2015. *Oscarella carmela* Muricy, Pearse, 2004. RBINS-IG 33439-POR.0127, Glutaraldehyde (this study), sea tables at Long Marine Lab. Santa Cruz, California, USA, S. Collected by Scott Nichols 01.08.2015. *Oscarella carmela* Muricy, Pearse, 2004, RBINS-IG 33439-POR.0128, Glutaraldehyde (this study), sea tables at Long Marine Lab. Santa Cruz, California, USA, S. Collected by Scott Nichols 01.08.2015.

#### Comparative material examined

Paratype B-6616 as *Oscarella carmela* specimen from an aquarium in J. Long Marine Laboratory. After examination it was find that this is *Oscarella pearsei* sp.nov. *Oscarella pearsei* Ereskovsky, Richter, Lavrov, Schippers, Nichols (this study). ZIN RAS № 11798, Ethanol. urn:lsid:zoobank.org:act:8431B41C-B08E-4205-8FE6-CCCFFB6EB574, sea tables at Long Marine Lab. Santa Cruz, California, USA, Collected by Scott Nichols 01.08.2015; ZIN RAS № 11800, Holotype, Glutaraldehyde. urn:lsid:zoobank.org:act:8431B41C-B08E-4205-8FE6-CCCFFB6EB574, sea tables at Long Marine Lab. Santa Cruz, California, USA, Collected by Scott Nichols 01.08.2015; ZIN RAS № 11799, paratype, Ethanol, sea tables at Long Marine Lab. Santa Cruz, California, USA, Collected by Scott Nichols 01.08.2015; ZIN RAS № 11801, paratype, Glutaraldehyde, sea tables at Long Marine Lab. Santa Cruz, California, USA, Collected by Scott Nichols 01.08.2015; RBINS-IG 33439-POR.0122, paratype, Ethanol, sea tables at Long Marine Lab. Santa Cruz, California, USA, Collected by Scott Nichols 01.08.2015; RBINS-IG 33439-POR.0123, paratype, Glutaraldehyde, sea tables at Long Marine Lab. Santa Cruz, California, USA, Collected by Scott Nichols 01.08.2015; RBINS-IG 33439- POR.0124 paratype, Glutaraldehyde, Ethanol, sea tables at Long Marine Lab. Santa Cruz, California, USA, Collected by Scott Nichols 01.08.2015. *Oscarella malakhovi* Ereskovsky, 2006 (ZIN RAS 10697 ZIN RAS 10698: Japan Sea) [28]. *Oscarella kamchatkensis* Ereskovsky, Sanamyan & Vishnyakov, 2009 (ZIN RAS 11058, ZIN RAS 11059 and ZIN RAS 11060: North Pacific, Avacha Gulf) [29]. *Oscarella lobularis* (Schmidt, 1862) and *Oscarella tuberculata* (Schmidt, 1868). NE Mediterranean Sea (Marseille region), underwater cave of Maire Island. *Oscarella microlobata* Muricy, Boury-Esnault, Bézac, Vacelet, 1996 and *Oscarella viridis* Muricy, Boury-Esnault, Bézac, Vacelet, 1996. NE Mediterranean Sea (Marseille region), underwater cave of Jarre Island [30]. *Oscarella balibaloi*, Pérez, Ivanišević, Dubois, Pedel, Thomas, Tokina, Ereskovsky, 2011. NE Mediterranean Sea (Marseille region), underwater cave of Maire Island [32]. *Oscarella nicolae* Gazave, Lavrov, Cabrol, Renard, Rocher, Vacelet, Adamska, Borchiellini, Ereskovsky, 2013 (MNHN DJV155, LSID urn:lsid:zoobank.org:act:DFFAD94B-9CAD-4F99-994D-BDEF75EF98A1, MNHN DJV156) North Sea, Norway (Bergen Fjords, Skarvoysundet) [25]. *Oscarella bergenensis* Gazave, Lavrov, Cabrol, Renard, Rocher, Vacelet, Adamska, Borchiellini, Ereskovsky, 2013 (MNHN DJV153, LSID:urn:lsid:zoobank.org:act:2D44BCFA-2163-47C7-9E70-EF6C13E0E4A4, MNHN DJV154) North Sea, Norway (Bergen Fjords, Skarvoysundet) [25].

#### Diagnosis (corrected from [1])

Known from the intertidal and subtidal, in and around Monterey Bay, California. Brown in color with bumpy, microlobate surface, wavy appearance, soft, slimy consistency. Mesohylar cells of two types with different granular inclusions, and vacuolar cells that also contain granular inclusions. Nucleus in choanocytes has basal position. Embryos follicle consists of one layer of flat cells. It has two morphotypes of endobiotic bacteria.

#### Description (corrected from [1])

Thinly encrusting, irregular sponges, generally light brown in color, up to 20–30 cm in diameter, with extremely soft, slimy consistency (Fig 1B and C). Thickness, color, and surface are variable. Most specimens <5 mm thick but sometimes ranging to ∼8mm. The surface is lumpy or wavy in appearance, microlobate with conspicuous channels and oscula. The sponge is easy to peel off smooth surfaces, as it attaches only at the tips of outgrowths that extend from the basopinacoderm.

#### Soft tissue organization

Spicule and fiber skeleton absent. Ectosome is 10 to 20 μm thick. Inhalant canals run perpendicular to the surface. Choanocyte chambers ovoid to spherical, eurypilous, from 45 to 65 μm in diameter, organized around large exhalant canals. Exhalant canals run towards a well-developed system of large basal cavities, separated by septa without choanocyte chambers.

### Cytology

**Exopinacocytes** (Fig 7A) flagellated with flat to oval shape from 9.5 to 14.3 μm wide by 2.8 μm high (N = 10). Cytoplasm includes electron translucent vacuoles 0.2 – 0.5 μm in diameter. Nucleus about 2.2 μm in diameter and basal in position. External surface of exopinacocytes covered by a layer of glycocalyx.

**Fig. 7.**
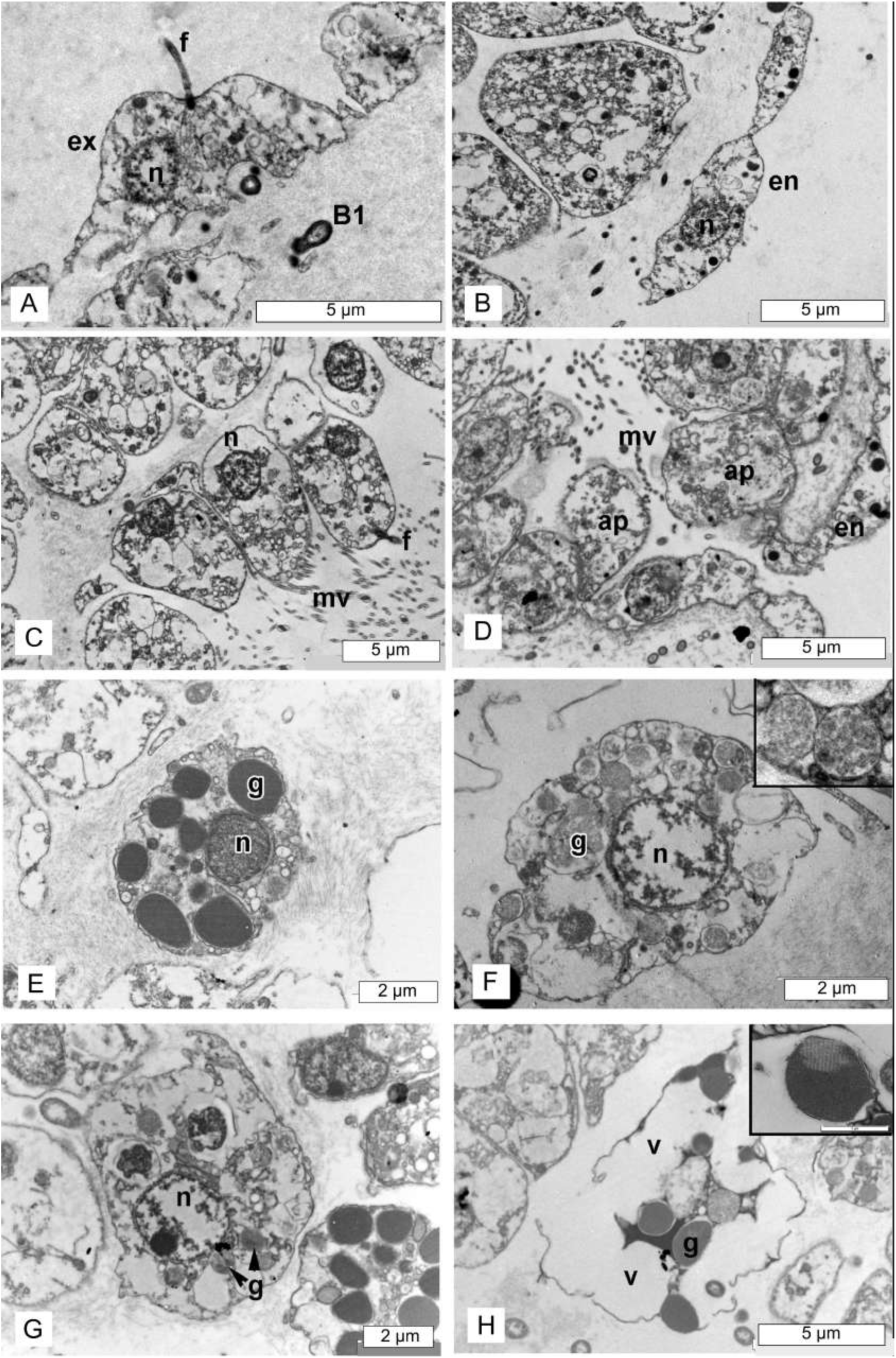
*Oscarella carmela* TEM of cells. **(A)** exopinacocyte; **(B)** endopinacocyte; **(C)** choanocytes; **(D)** apopylar cell; **(E)** granular cell type 1; (**F)** differentiated granular cell type 2, Inset: granules with filamentous inclusions; **(G)** granular cell type 2 with small number of inclusions; **(H)** vacuolar cell with granular inclusions, Inset - granule. (Abbreviations: ap – apopylar cell, B1 – bacteria type 1, en – endopinacocyte, ex – exopinacocyte, f – flagellum, g – granule, mv – microvilli, n – nucleus, v – vacuole).

**Endopinacocytes** (Fig 7B) flagellated, from 8.3 to 13.7 μm wide by 2.9 μm high (N = 10). Nucleus ovoid (1.8 x 3.1 μm in diameter). Cytoplasm includes electron dense granules 0.2 - 0.8 μm in diameter.

**Choanocytes** (Fig 7C) ovoid to trapezoidal, about 4.4 μm wide by 8.2 μm high (N = 20). Nuclei (∼2.1 μm diameter) have basal position without any contact with flagellar basal apparatus. Cytoplasm includes electron transparent vacuoles from 0.5 to 1.9 μm in diameter.

**Apopylar cells** (Fig 7D) roughly triangular or oval in section, 6.3 μm wide by 4.7 μm high (N = 5). Nucleus spherical, up to 2.9 μm in diameter. Cytoplasm contains mitochondria, digestive vacuoles, and small osmiophilic inclusions.

Surface of endopinacocytes, choanocytes and apopylar cells covered by thin irregular layer of glycocalyx. Choanoderm and pinacoderm underlined by continuous basement membrane 0.9 - 1.7 μm thick.

#### Three types of cells with inclusions occur within the mesohyl

**Granular cells type 1** (Fig 7E) (“type 1 cell with inclusions” described by [1]) are ovoid, 4.5 x 7 μm, irregular, with short pseudopodia. Nucleus about 1.9 μm in diameter, ovoid or compressed by the abundant cytoplasmic inclusions. Cytoplasm filled with oval inclusions of one kind, 0.8–1.9 μm wide, with electron dense and homogeneous contents.

**Granular cells type 2** (Fig. 7F, G) (“archeocytes”, described by [1]). Irregular to ovoid 5.5 x 7.9 μm, sometimes with short pseudopodia. Cytoplasm filled with abundant spherical inclusions with filamentous contents 0.7 μm in diameter and with some phagosomes 0.9 to 1.7 μm in diameter. It appears that granules fuse with each other, resulting in the formation of electron transparent vacuoles from 0.7 to 2.5 μm in diameter. Nucleus is ovoid or irregular, 2.4 μm and sometimes a nucleolus is evident.

**Vacuolar cells with granular inclusions** (Fig 7H) (type 2 cell with inclusions (“spherulous vacuolar cell”) described by [1]) have irregular shape, 7.1 x 13.0 μm. Cytoplasm includes 4 - 15, 0.8 – 1.8 μm oval inclusions with homogenous osmiophilic contents, however central, or lateral part of this inclusion are clear (light-gray). There are 3 - 7 larger vacuoles from 2.1 to 7.1 μm in diameter, with clear and filamentous contents. Nucleus 1.6 – 1.9 μm in diameter without nucleolus, ovoid or compressed by abundant cytoplasmic inclusions. This cell type may be involved in the secretion of extracellular matrix, because at the end of the ontogeny of these cells, vacuoles appear to fuse into one or two large vacuoles, their membrane seems to merge with cell membrane and vacuolar content is presumably released in the mesohyl (Fig 7H). This appears to be the maturation sequence of granules (S4 Fig).

#### Two morphological types of extracellular, endobiotic bacteria occur in the mesohyl

Type B1 (Fig 8A and B) are abundant, ovoid 1.1 - 1.5 μm in length and 0.7 μm in diameter (N = 22). Cell wall consists of two membranes. Cytoplasm filled with loose filamentous materials concentrated between the cell wall and central clear part (0.17 μm). Nucleoid is loose filamentous network and with thin filaments closer to the periphery. Surface is even or slightly wavy. Type B2 (Fig 8C and D) is rod-like, about 1.1 μm length and about 0.24 μm in diameter. Cell wall consists of two membranes. Cytoplasm filled with dark filamentous materials, and without a clear distinction between nuclear and cytoplasmic regions.

**Fig. 8.**
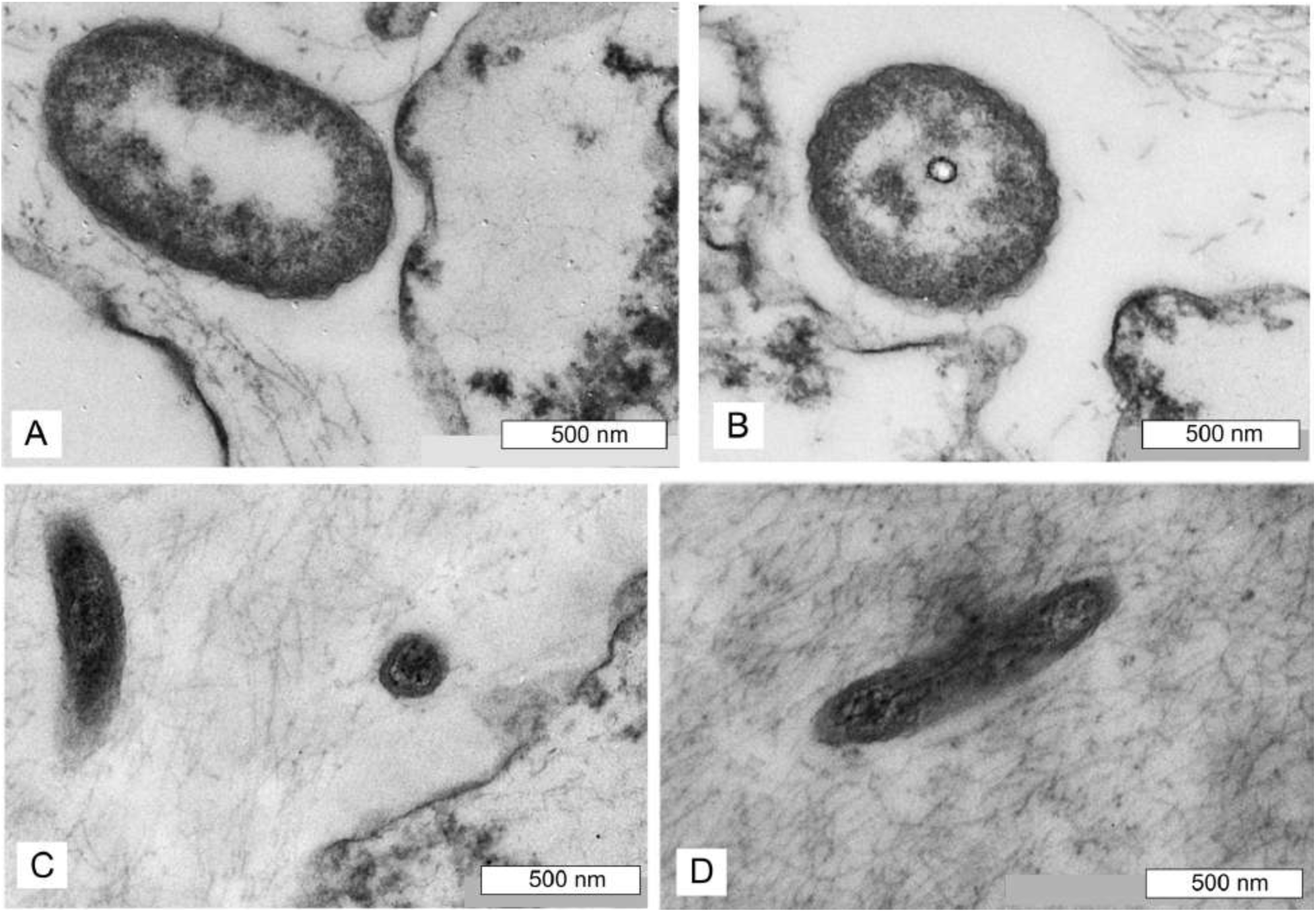
*Oscarella carmela* TEM of symbiotic bacteria. **(A, B)** symbiotic bacteria type 1; **(C, D)** symbiotic bacteria type 2.

#### Reproduction

*Oscarella carmela* is viviparous and simultaneously hermaphroditic: male and female reproductive elements are present in the same individuals (Fig 9A). Oogenesis and embryogenesis are asynchronous; all stages from oogonia to egg were observed within the same specimen. Mature eggs are isolecithal and polylecithal, with a cytoplasm full of yolk granules with diameter about 48-58 μm (Fig 9A). Embryogenesis is also asynchronous, resulting in formation of cinctoblastula larvae (Fig 9D). Eggs and embryos (about 80 μm in diameter) are located in the basal part of the choanosome and are surrounded by a follicle (Fig 9 A-D). Follicle is simple from oocyte vitellogenesis stage (Fig 9A and E) to embryos and larvae (Fig 9B-D and F) and consists of only one layer of flat cells. The spermatic cysts are oval and have different dimension (from 28-52 x 21-48 μm) and are randomly distributed in the sponge mesohyl (Fig 9 G). Spermatogenesis is generally asynchronous inside spermatic cysts (Fig 9G and H).

**Fig. 9.**
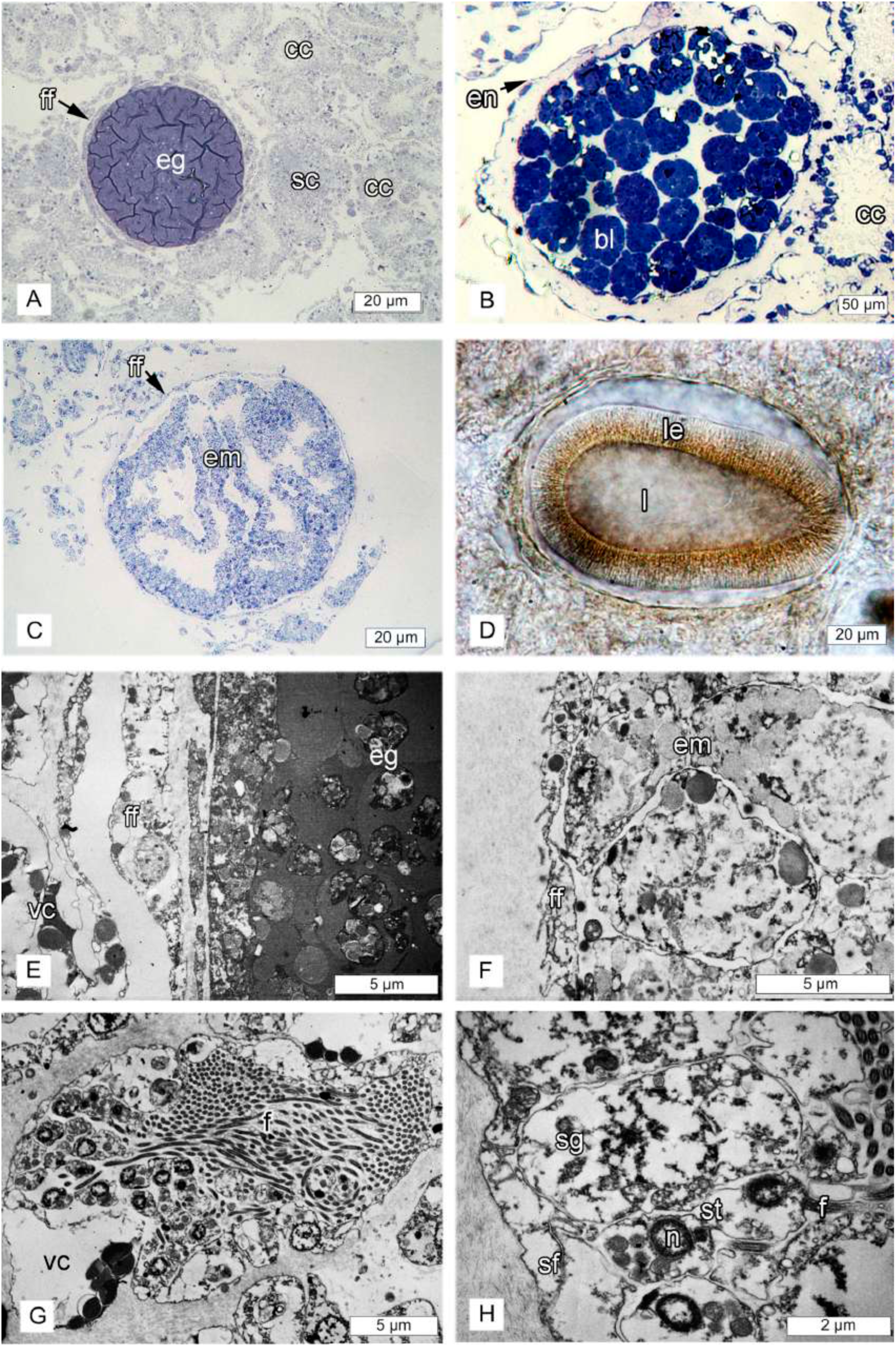
*Oscarella carmela* follicle and reproduction. **(A)** semi-thin section of the egg with flat monolayer follicle; **(B)** semi-thin section of cleaving embryo with flat monolayer follicle; **(C)** semi-thin section of the embryo with flat monolayer follicle; **(D)** *in vivo* image of cinctoblastula larva; **(E)** TEM of flat monolayer follicle of the egg; **(F)** TEM of with flat monolayer follicle of the embryo; **(G)** spermatocyst; **(H)** detail of the spermatocyst with surrounding follicle epithelium, spermatogonia and the spermatids. (Abbreviations: B1 – bacteria type 1, B2 – bacteria type 2, bl – blastomeres; cc - choanocyte chamber, eg – egg, em – embryon, f – flagella, ff – flat follicle, l – larva, le – larval epithelium, n – nucleus, sf - follicle of a spermatocyst, sg - spermatogonia, st – spermatid, vc – vacuolar cell with granular inclusions).

### Taxonomic remarks

The absence of spherulous cells with paracrystalline inclusions is consistent with the phylogenetic placement of *Oscarella carmela* in the clade that includes *O. lobularis, O. tuberculata, O. viridis, O. rubra, O. malakhovi, O. bergenensis* [1, 25, 28, 30, 32]. The slimy consistency and abundant mucus of *Oscarella carmela* resembles *O. nicolae* [25] and *O. balibaloi* [31] but differs significantly from *O. pearsei* sp. nov.

## Discussion

Prior to this study there were 19 valid species of the genus *Oscarella* described, worldwide [33]. Only five of these were found in the Pacific: *O. nigraviolacea, O. stillans* [34] in the Southern Hemisphere and *O. carmela* [1], *O. malakhovi* [28] and *O. kamtchakensis* [35] in the Northern Hemisphere. Characteristics of *Oscarella* that are observable in the field, such as coloration, surface morphology and consistency are difficult to use for accurate species discrimination because they can vary within a single species. For example, in the Mediterranean, *O. lobularis* and *O. tuberculata* may appear in several different colors: white, yellow, green, red, violet, blue [32, 35, 36]. Likewise, *O. balibaloi* [31] varies in color from white to yellow, orange and violet. Characteristics such as tissue consistency and surface morphology are potentially less variable within each species, but they are also more subjective to describe.

The initial clue that the species *Oscarella carmela* may represent two distinct species was that the EST dataset published under the name *O. carmela* [2] was significantly divergent from the Illumina transcriptome that was later sequenced from a separate individual (this study). Here, we correct the species attribution of the published ESTs and draft genome from *O. carmela* to *O. pearsei* sp. nov., and we report correctly attributed Illumina-sequenced transcriptomes for each of these species. The mitochondrial genome previously attributed to *O. carmela* [3] remains unchanged.

*O. carmela* and *O. pearsei* sp. nov. are co-distributed along the central California coast, albeit their distributions are incompletely known. We sampled tissue from individuals of each species (verified both morphologically and by DNA barcoding), growing side by side on the same rock at the *O. carmela* type locality in Carmel, California. We also found *O. carmela*, but not *O. pearsei* sp. nov., in the shallow subtidal zone of Pacific Grove, California on the underside of rocks. Both species can be found growing in sea tables at the Joseph Long Marine Laboratory, University of California Santa Cruz. Specimens of *Oscarella* have also been found growing in the intertidal in the Santa Barbara region to the south, and in Coos Bay Oregon to the north (Jeff Goddard, pers. comm.). The identity of these species is unknown, but we obtained a sample from Santa Barbara (gift from Desmond Ramirez to K.S.) and performed DNA barcoding and found it to represent a third species, distinct from either *O. carmela* or *O. pearsei* sp. nov. It is possible that multiple, undescribed species exist in the Eastern Pacific. This situation resembles the case of *Oscarella* systematics in the Mediterranean, where a single species, *O. lobularis*, was initially recognized but eventually many co-distributed species were described on the basis of their cytological differences and later, using DNA evidence (see [37] [35]).

*Oscarella pearsei* sp. nov is readily distinguished from *O. carmela* using LSU and mt-cob barcodes, and phylogenetic analyses of SSU, LSU and mitochondrial genomic data.

Interestingly, these two species represent two major branches in the family Oscarellidae and are only distantly related to each other. Furthermore, *O. pearsei* sp. nov is currently the only species in the family with a distinct mitochondrial gene order, as a result of an inversion encompassing three protein-coding and four tRNA genes.

In addition to genetic evidence, careful re-analysis of the morphology of *O. carmela* and *O. pearsei* sp. nov. reveals characteristics useful for identification in the field and in the lab. With respect to gross morphology, *O. pearsei* sp. nov. has a light tan to orange color while *O. carmela* is almost exclusively light-brown. Additionally, the surface of *O. pearsei* sp. nov. is finely and uniformly lobate with small hemispherical lobes that are unique for the genus. In contrast, the surface of *O. carmela* is lumpy or wavy appearance and microlobate; similar to *O. microlobata, O. kamchatkensis, O. malakhovi*. and *O. balibaloi* [28, 30, 31, 35]. Both *O. carmela* and *O. pearsei* sp. nov. are soft and delicate to the touch, but *O. carmela* is generally thicker, the less delicate of the two, and produces much more mucus.

Because *Oscarella* lacks skeletal elements (inorganic and organic) the best morphological characters for reliable species discrimination are ultrastructural: each species has unique cell type combinations in the mesohyl, and unique and consistently associated bacterial morphotypes. Specifically, *O. pearsei* sp. nov. differs from all other described *Oscarella* species, including *O. carmela* (Table 3) in that it lacks vacuolar cells, but contains archeocytes, abundant granular cells of a single type, and spherulous cells with paracrystalline inclusions. In contrast, *O. carmela* contains vacuolar cells, has two discrete types of granular cells and lacks both archeocytes and spherulous cells with paracrystalline inclusions. The latter cell type is a synapomorphy for the clade to which *O. pearsei* sp. nov. belongs, and unites this species with *O. microlobata, O. imperialis, O. kamchatkensis, O. balibaloi, O. nicolae* and *Pseudocorticium jarrei* [25, 30, 31, 35, 38].

**Table 3.**
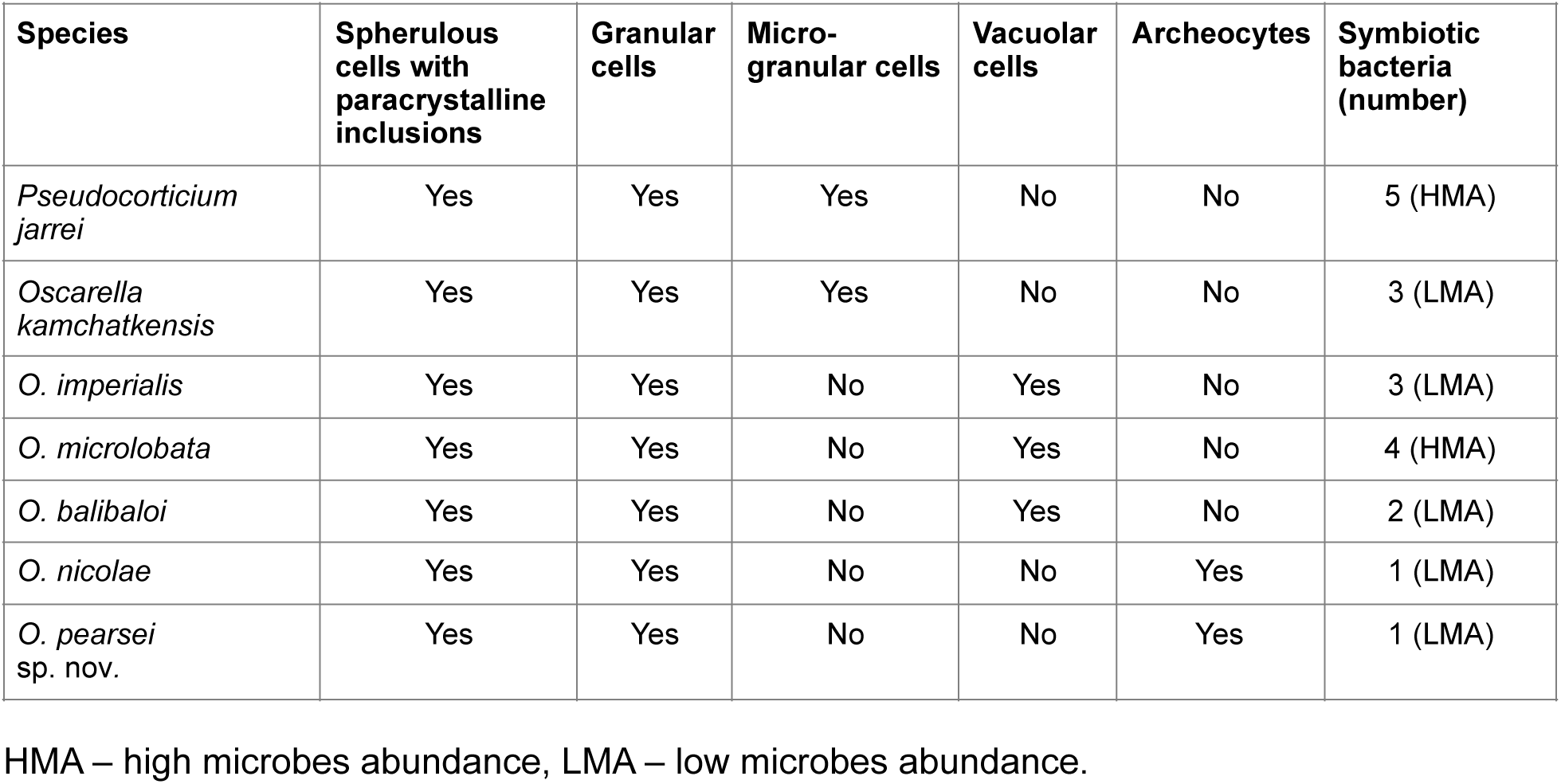
The cytological characters of Oscarellidae species with cells that contain paracrystalline inclusions.

The granular cells of *O. pearsei* sp. nov. also differ from granular cells of other *Oscarella* species due to the presence of an unusually osmiophilic central region within vacuoles, which is surrounded by opaque (gray) peripheral content. In comparison, in *O. balibaloi*, granular cells have entire homogeneous, electron-dense inclusions with a crystalline texture that is visible under high magnification [31]. In *O. nicolae* and *O. kamchatkensis* granular cells, the granules are also entirely homogeneous and electron dense [25, 29]. In *O. microlobata* granular cells (type I), the cytoplasm is almost completely filled with heterogeneous inclusions, containing 10 - 30 osmiophilic granules embedded in a dense matrix [30]. Finally, granular cells (type 1) of *O. imperialis* are almost completely filled by 2 – 5 μm spherical vacuoles, and 5 −20 μm partly paracrystalline inclusions (i.e., the inclusions have a partly paracrystalline organization and are partly amorphous) [30].

The final cytological characteristic that is useful for distinguishing between *O. carmela* and *O. pearsei* sp. nov. is the position of the nucleus in choanocytes. We find that, with absolute consistency, choanocytes in *O. pearsei* sp. nov. have an apically positioned nucleus (just below the microvillar collar) whereas choanocytes in *O. carmela* have a basally positioned nucleus.

A limitation of using cytological characteristics for species discrimination is that it is labor intensive. For the most part, tissue has to be prepared for examination by transmission electron microscopy. Fortunately, in reproductive individuals of *O. carmela* and *O. pearsei* sp. nov. it is also possible to consider a unique histological characteristic: the morphology of the embryonic follicle epithelium. This is not a traditional character considered in the early taxonomic literature for *Oscarella*, but we find that it reliably distinguishes between these species and that it is even visible in thin, living samples and tissue ‘squashes’ when viewed with phase microscopy. Specifically, *O. carmela*, has flat monolayer follicle epithelium, whereas the follicle epithelium of *O. pearsei* sp. nov. is prominent and composed of cuboidal epithelial cells which persist from the stage of vitellogenesis to the stage of larval release. The presence of a cuboidal follicle epithelium has previously been described only from *O. nicolae* [25], which is found in the North sea, Norway, and cannot therefore be confused with *O. pearsei* sp. nov.

Based upon this study, we caution that previously published references to the species *O. carmela* were incorrect, and that the actual species under study was what we have now described as *O. pearsei* sp. nov. The only exception to this is the published mitochondrial genome for *O. carmela*, and studies that reference those data. Henceforth, we have corrected the species names associated with all publicly available genetic resources (i.e., ESTs, transcriptomes, genomes, and mitochondrial genomes), including in the NCBI databases, and compagen.org [23]. Future studies that involve new collection of *Oscarella* tissue from the Central California coast (or elsewhere) should be diligent to keep samples individually separated and to save voucher material for DNA barcoding and possibly morphological analysis.

## Acknowledgements

S.A.N. would like to acknowledge Nicole King at the University of California Berkeley, where some of this research was conducted, and Betsy Steele and John Pearse for facilitating access to Long Marine Laboratory and for help with field collections. Thanks to Guilherme R. S. Muricy from Museu Nacional Rio de Janeiro (MNRJ) for the opportunity to work with slides of the type material. D.V.L. thanks Walker Pett (Iowa State University) for his help with the analysis of genomic data. A.E. gratefully thanks Joël Courageot and Alexandre Altié of Service Commun de Microscopie Électronique et Photographie Faculté de Médecine La Timone, Aix-Marseille Université and Daria Tokina and Sandrine Chenesseau for technical support.

---

**S1 Fig.**
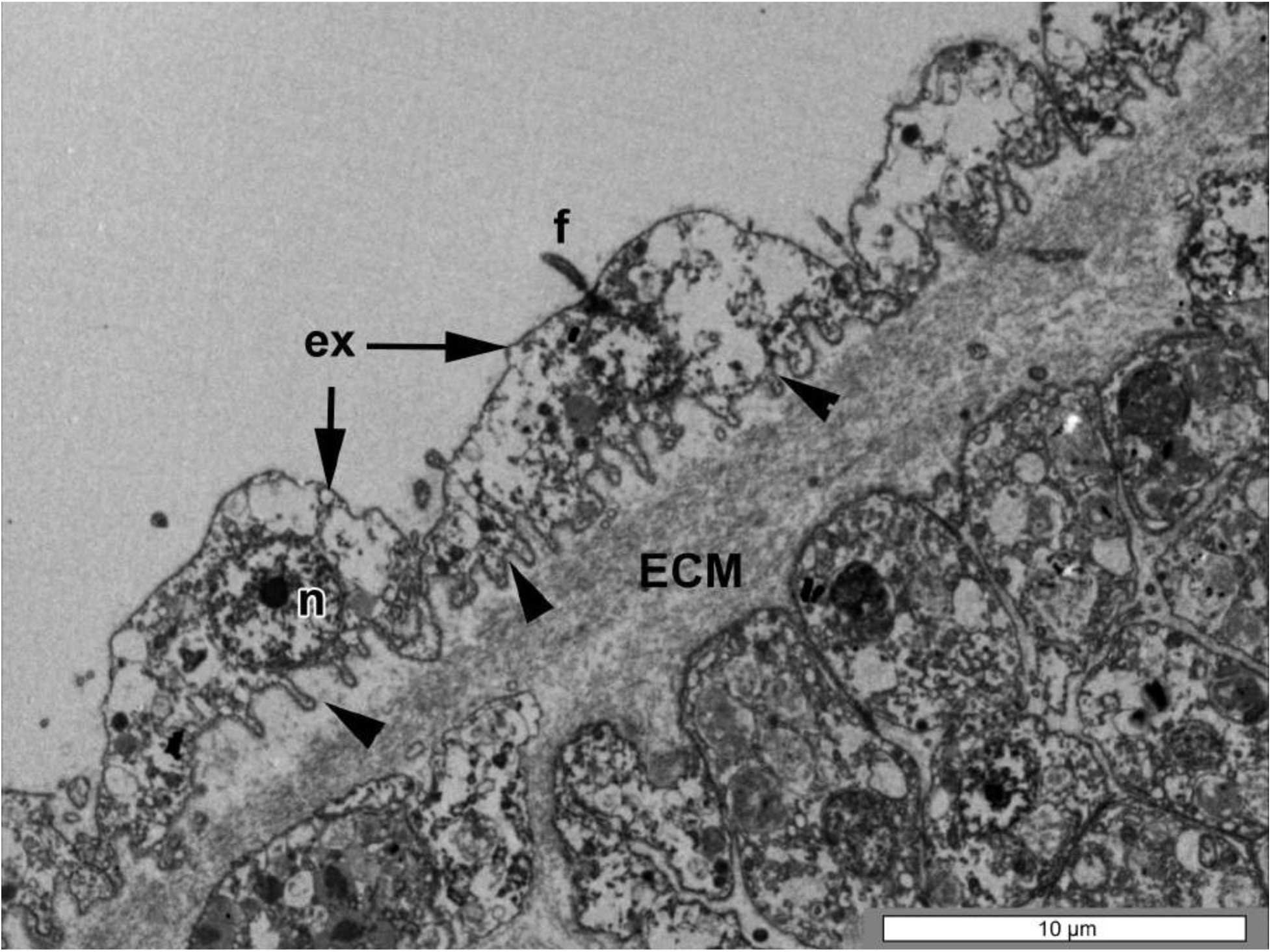
TEM of an exopinacocyte (ex) of *Oscarella pearsei* sp. nov., showed basal surface with filopodia (arrowheads) that extend into the extracellular matrix (ECM). f – flagellum, n – nucleus.

**S2 Fig.**
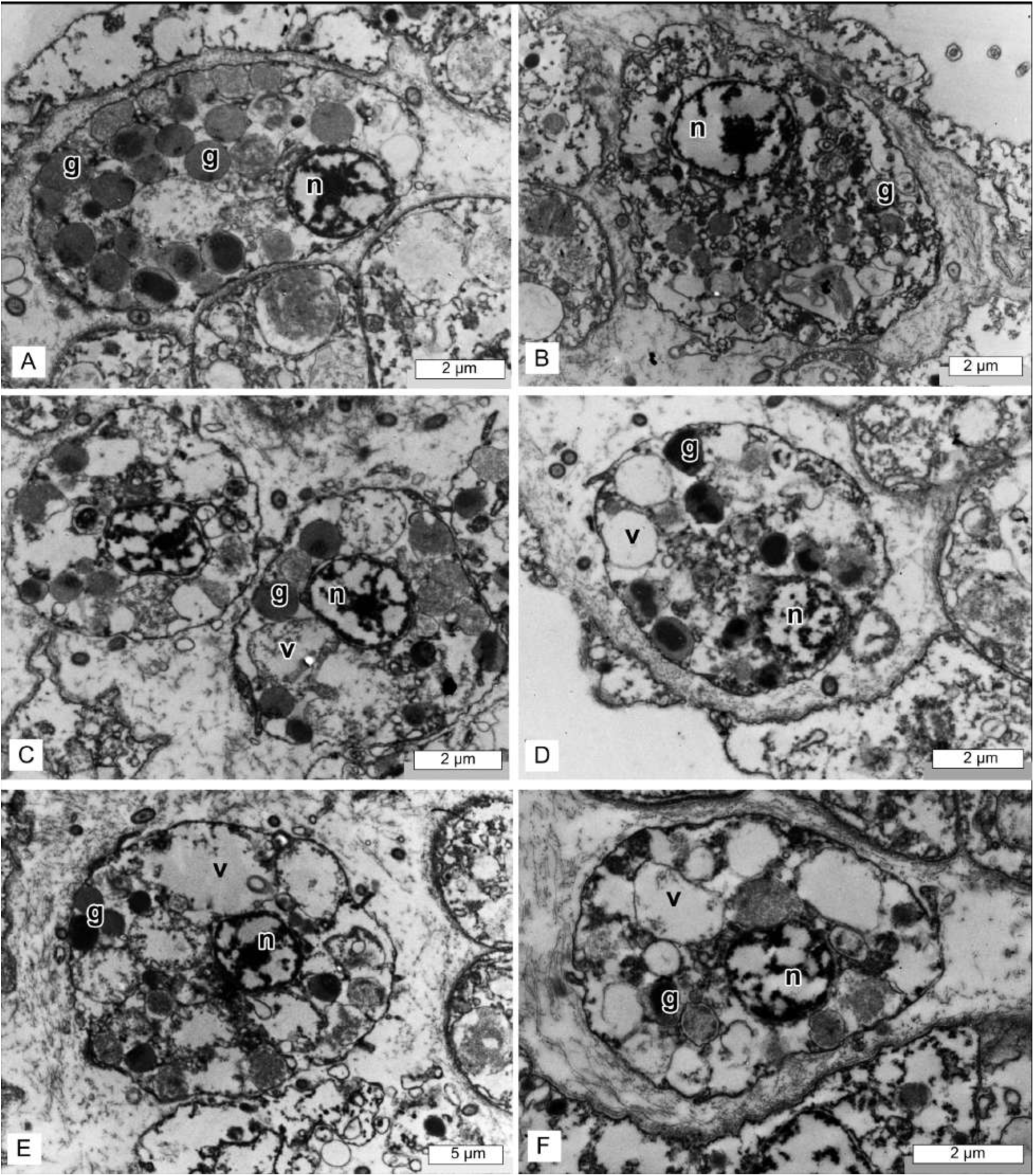
*Oscarella pearsei* sp. nov. TEM of granular cells at different stage of their ontogeny. g – granule, n – nucleus, v – vacuole.

**S3 Fig.**
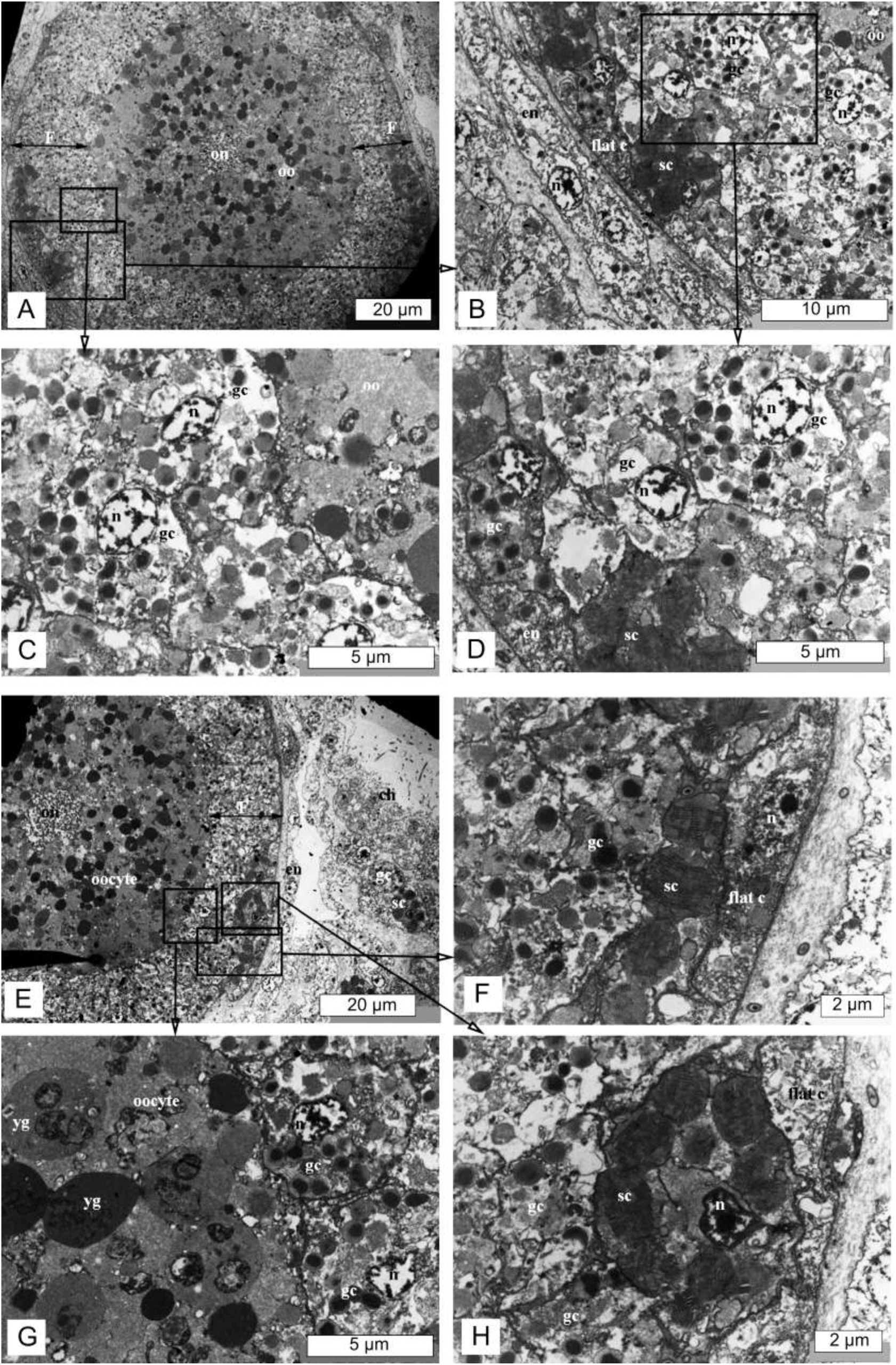
*Oscarella pearsei* sp. nov. TEM of follicle surrounding mature oocyte. **(A)** Mature oocyte with two-layer follicle. **(B) - (D)** details of **(A)** showing granular and spherulous cells, composed internal layer of the follicle. **(E)** the part of a mature oocyte with follicle. **(E) - (H)** details of **(E)** showing granular and spherulous cells, composed internal layer of the follicle and external flat cells of a follicle **(F), (H)**. ch – choanocytes, en – endopinacocytes, F – follicle, flat.c – flat cells of follicle, gc – granular cell, on – oocyte’s nucleus, oo – oocyte, n - nucleus, sc- spherulous cells with paracrystalline inclusions, yg – yolk granules.

**S4 Fig.**
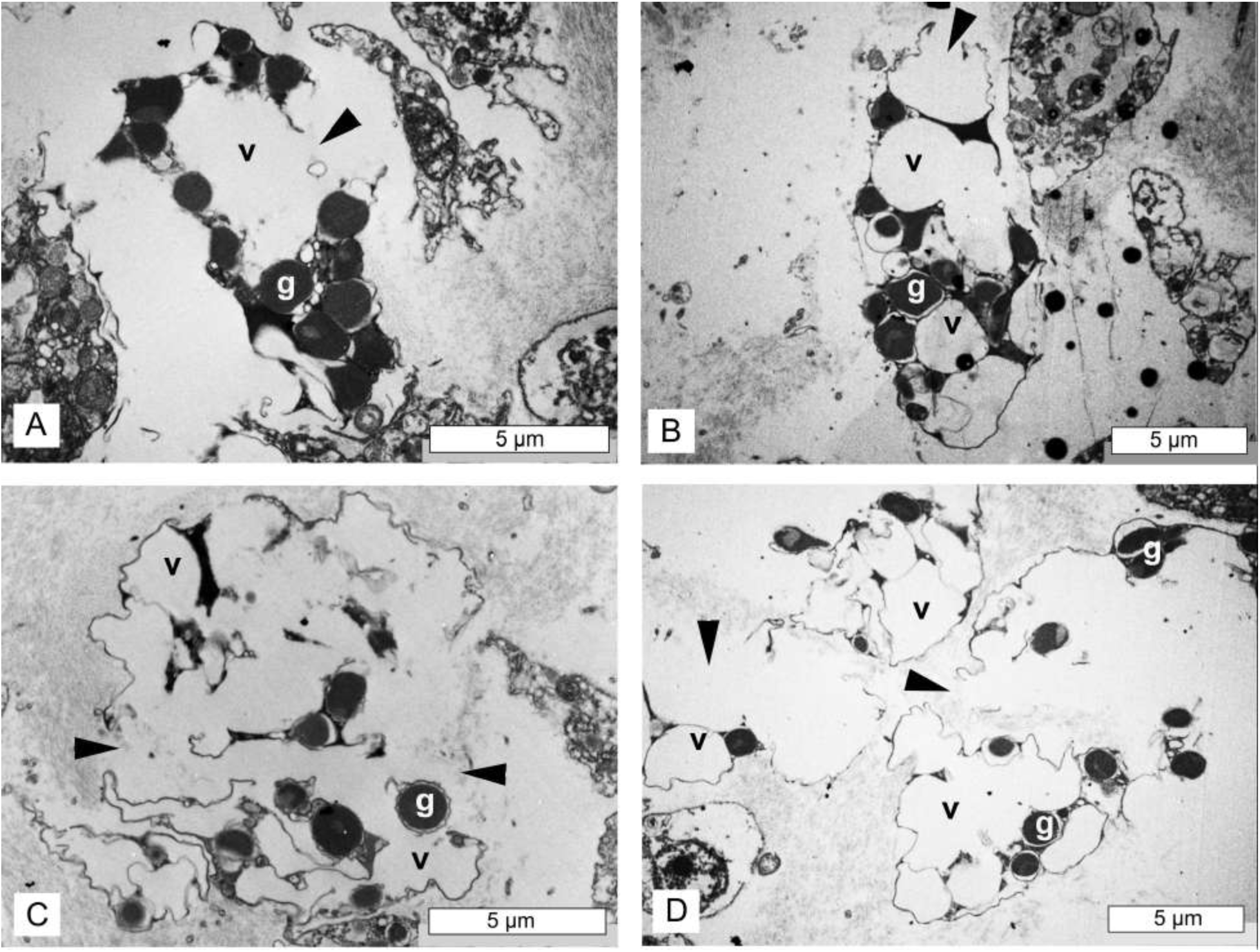
*Oscarella carmela* TEM of vacuolar cells with granular inclusions at different stages of their ontogeny. Arrowheads show the apparent release of vacuolar content in the mesohyl. g – granule, v – vacuole.

